# Shape, Strain, and Stability: Epithelia Under High Strain

**DOI:** 10.1101/2025.11.12.687918

**Authors:** Jing Yang, Yicheng Dong, Carter B. Jones, Yingli Wang, Chloe V. Merino, Carsten Stuckenholz, Lance A. Davidson

## Abstract

Mechanical forces play crucial roles in epithelia tissue morphogenesis and function, yet individual cells often respond differently to the same external force. Strain heterogeneity, the non-uniform deformation of cells under uniform external mechanical stimuli, may underlie tissue-level robustness and guide morphogenetic outcomes. In this study, we applied a defined uniaxial strain to *Xenopus laevis* epithelial explants and investigated how strain was distributed at the cellular level. Live imaging and quantitative analysis revealed that despite a uniform tissue-level strain, cellular strain was heterogeneous, suggesting variable responses to a uniform mechanical stimulus. To understand the source of this variability, we used an image analysis pipeline to explore multiple aspects of cell morphology. We found that cell intrinsic material properties, represented by Poisson index, had the strongest correlation with strain heterogeneity, suggesting a dominant role in variable mechanical response. We further analyzed how force was distributed at the cellular level using a vinculin force sensor and laser ablation. These experiments demonstrated that forces are primarily transmitted through the medio-apical actomyosin cortex whereas junctional actomyosin facilitates dissipation and remodeling. These findings provide new insights into the physical principles that underlie epithelial resilience and adaptive remodeling, highlighting the importance of distinct functions of junctional and medio-apical actomyosin networks in mechanical adaptation.

**Graphical Abstract:** 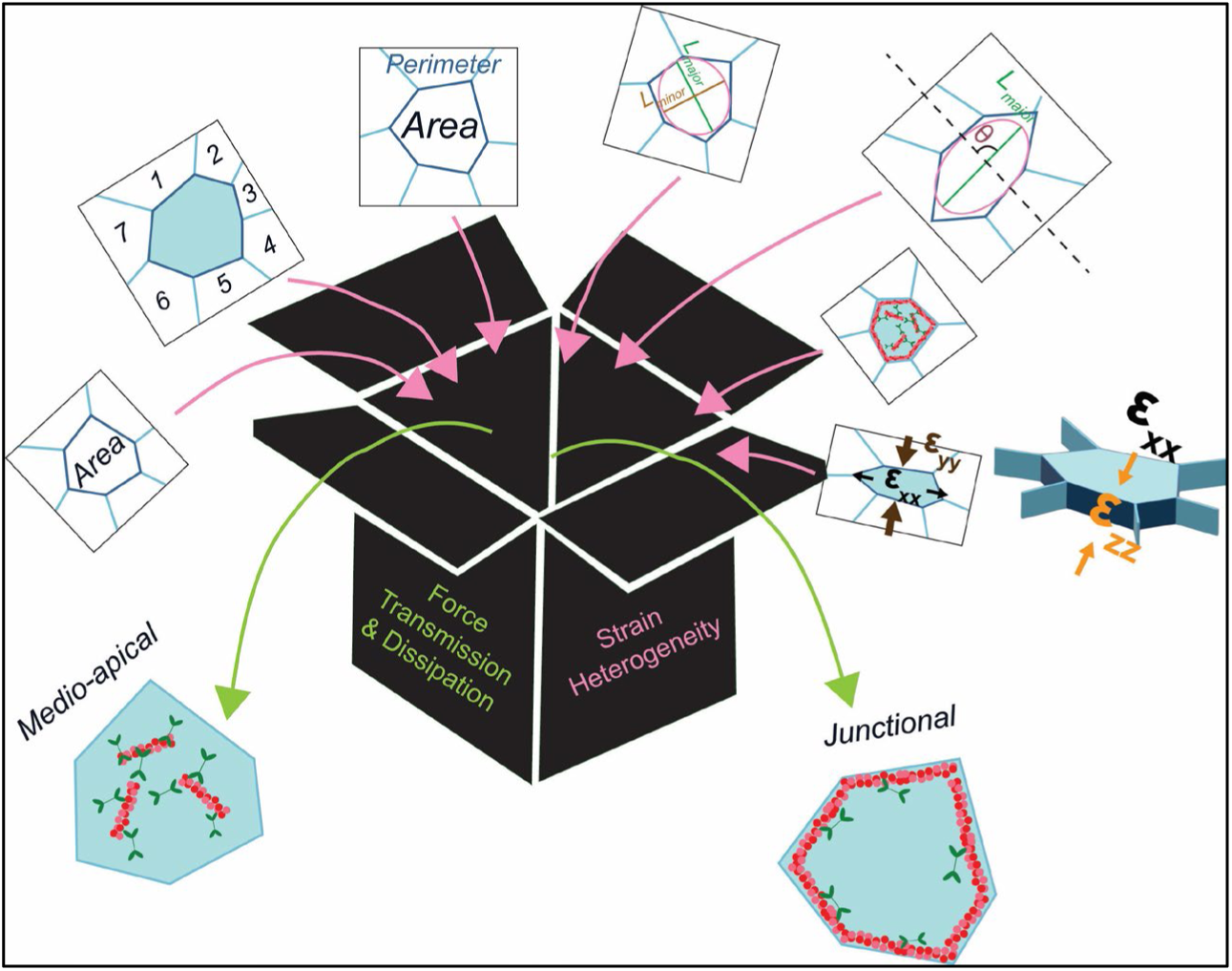

## 1. Introduction

Mechanical forces play crucial roles in metazoans, influencing processes ranging from embryonic development to tissue homeostasis and disease progression (Ingber et al., 1995; Engler et al., 2006; Davidson et al., 2009; Gjorevski and Nelson, 2010; Feroze et al., 2015; Heer and Martin, 2017; Northcott et al., 2018; Kim et al., 2020). Mechanical forces shape vertebrate embryos at early stages by regulating cellular behaviors (Thompson et al., 2019; Chu et al., 2020). For example, mechanics can drive stem cell differentiation (Engler et al., 2006), guide neural crest migration during development (Barriga et al., 2018; Shellard and Mayor, 2021), or trigger a mesenchymal-to-epithelial transition in heart progenitor cells (Jackson et al., 2017; Kim et al., 2020). In epithelial tissues, mechanical forces regulate epithelial morphogenesis (Arnold et al., 2019), homeostasis (Anjum et al., 2024; Tarannum et al., 2024; Landino et al., 2025), tissue integrity (Nestor-Bergmann et al., 2022), and tissue function (Chien et al., 2015; Nestor-Bergmann et al., 2019). Within tissues, mechanical forces are transmitted through complex pathways that involve cell-cell adhesions (Arnold et al., 2019; Manning et al., 2019), the extracellular matrix (ECM) (Conway et al., 2023), and intracellular cytoskeletal networks (Shyer et al., 2017). Junctional complexes and intracellular cytoskeleton facilitate the transfer of forces between neighboring cells (Gomez et al., 2011; Vasquez and Martin, 2016), while the ECM provides a structural framework that also transmits external forces across the tissue (Maller et al., 2010; Bachir et al., 2017). At the cellular level, cytoskeletal components such as actomyosin networks enable cells to sense, resist, and adapt to mechanical forces (Vasquez and Martin, 2016; Bachir et al., 2017; Vijayraghavan and Davidson, 2017).

Key to the mechanical roles of epithelia during morphogenesis is the ability to rearrange relative to one another while maintaining structural integrity (Jung et al., 2006; Heer and Martin, 2017). This ability to rearrange is commonly referred to as tissue fluidity or by describing the tissue as “fluid-like” (Guillot and Lecuit, 2013; Bi et al., 2016). Like a nonbiological material that can change material properties under mechanical strain, tissue properties can oftentimes change under large mechanical strain by inducing transitions between solid-like and fluid-like states (Guillot and Lecuit, 2013; Bi et al., 2016; Banavar et al., 2021). This occurs through “jamming” and “unjamming” mechanisms, where high strain either increases tissue rigidity (jamming) or enhances cellular mobility and rearrangement (unjamming), fluidizing intracellular cytoskeleton (Bi et al., 2016; Atia et al., 2021; Blauth et al., 2021). Several factors influence solid-fluid transitions in epithelial cells, including cellular adhesion (Kowalczyk and Green, 2013), cytoskeletal organization (Guillot and Lecuit, 2013), and ECM composition (Kozyrina et al., 2020; Banavar et al., 2021; Palmquist et al., 2022). For example, strong cell-cell adhesion or feedback systems may reduce fluidity, while a more dynamic cytoskeleton network can enhance it (Clarke and Martin, 2021; Founounou et al., 2021). The cell shape index reports cell geometry and has been correlated with tissue fluidity (Bi et al., 2016). Cells having a larger shape index, e.g., less circular, are suggested to be more fluid-like, reflecting higher levels of cell-cell neighbor exchange (Bi et al., 2016; Erdemci-Tandogan et al., 2018; Jain et al., 2020).

Due to the complexity of mechanosensing and mechanotransduction, individual cells within an epithelial tissue may exhibit different mechanical strain when under the same level of force (Gjorevski and Nelson, 2012; Li et al., 2019). For example, while one cell might elongate, a neighboring cell may undergo minimal deformation. It has been suggested that such heterogeneity in cellular strain can affect downstream processes, including cell signaling, differentiation, and tissue remodeling (Tsuboi et al., 2017; Li et al., 2019). In epithelial tissues, the distribution of heterogeneous strain can create directional differences in mechanical properties or force transmission, leading to anisotropic mechanical cues that regulate organ-level functions by guiding cell alignment, migration, and signaling (Sugimura and Ishihara, 2013; Tsuboi et al., 2017; Huebner et al., 2021). Understanding the origin of heterogeneity is key to gaining insights into tissue mechanics and functionality; however, the mechanisms responsible for such heterogeneity remain unknown.

Several factors may contribute to the variability in how epithelial cells respond to mechanical strain. First, cells can vary in stiffness; therefore when under the same forces, cells may exhibit different mechanical strains, i.e., stiffer cells exhibit lower strain while softer cells exhibit higher strain (Li et al., 2016; Bodenschatz et al., 2022). Cell geometry and topology may also affect cellular strain heterogeneity. Differences in cell shape, size, and arrangement within a tissue can lead to variations in how mechanical forces are distributed and transmitted (McBeath et al., 2004; Lecuit and Lenne, 2007; Oakes et al., 2014). For example, hexagonal packing in epithelial layers optimizes force transmission and minimizes strain concentrations (Gibson et al., 2006; Sugimura and Ishihara, 2013; Eckert et al., 2023). Additionally, the local microenvironment, including variations in ECM composition and cell-cell adhesion, may modulate local cell responses to mechanical cues (Zhang et al., 2009; Zhou et al., 2009; Bachir et al., 2017; Holle et al., 2018; Sumi et al., 2018; Arnold et al., 2019). Cells on dense ECM may experience different mechanical forces compared to cells in more compliant environments (Saraswathibhatla et al., 2023; Crossley et al., 2024). Intrinsic cellular properties, such as cytoskeletal organization or cytoarchitecture, may also contribute to heterogeneous responses (Luby-Phelps, 2000). Variations in the density or activity of actomyosin networks or intermediate filament density can alter how cells deform and transmit forces, influencing their ability to adapt to mechanical stress (Munjal and Lecuit, 2014; van Bodegraven and Etienne-Manneville, 2021; Ndiaye et al., 2022). Additionally, substructural alignment and orientation of the cytoskeletal components within cells can generate an anisotropic mechanical response (Chanet et al., 2017). While intrinsic factors such as variations in cellular stiffness, geometry, and cytoskeleton organization contribute to this heterogeneity, some degree of variability in protein abundance or activity may also arise from stochastic noise. Lastly, cells continuously experience intrinsic and extrinsic forces that can introduce random fluctuations in their shape, and thus strain (Guo et al., 2014; Chang and Marshall, 2017).

Actomyosin networks are thought to be a primary driver of force generation, transmission, and dissipation within embryonic cells and tissues (Munjal and Lecuit, 2014; Chanet et al., 2017). These networks generate force by myosin II contraction, which pull on actin filaments to create tension that regulates cells shape, adhesion, and mechanical interactions within tissues (Chanet et al., 2017; Arnold et al., 2019; Kim et al., 2020). Actomyosin networks play a crucial role in force transmission by linking to cell-cell junctions and the extracellular matrix, enabling coordinated mechanical responses across tissues. Cells can dynamically respond to mechanical cues by remodeling, recruiting or disassembling actomyosin components (Munjal and Lecuit, 2014; Vasquez and Martin, 2016; Chanet et al., 2017; Jackson et al., 2017). This adaptive capability allows epithelial cells to modulate their mechanical properties, facilitating tissue remodeling and enabling transitions between different mechanical states. This adaptability is especially evident in distinct subcellular regions, such as junctional, medio-apical, and basolateral domains, where actomyosin plays specialized roles in force transmission and dissipation (Shawky and Davidson, 2015; Arnold et al., 2019; Rosa-Birriel et al., 2024). Junctional actomyosin, which localizes next to cell junctions, is crucial for maintaining epithelial integrity and coordinating mechanical signals across the tissue (Quiros and Nusrat, 2014; Arnold et al., 2019). Medio-apical actomyosin, which assembles at the apical cortex, is involved in regulating apical tension and modulating epithelial shape change during morphogenesis (Arnold et al., 2019; Rosa-Birriel et al., 2024). Basolateral actomyosin is essential for collective migration by generating pulsatile contractions through focal adhesions and regulating tissue stiffness during development through cortex thickness (Hidalgo-Carcedo et al., 2011; Shawky et al., 2018). In each of these roles, actomyosin contractility maintains force balance in the tissue and underlies the timely response to mechanical stress; however, the relative contributions of different actomyosin networks to force transmission and dissipation remain unexplored.

Multiple sources of heterogeneous mechanics of cells and their roles in generating, transmitting, and dissipating forces emerge from observations of embryonic morphogenesis, tissue remodeling, and organoid self-assembly (Tsuboi et al., 2017; Huebner et al., 2021). By necessity, these observations coincide with force generation by the same tissues that self-assemble into a mechanical structure. Efforts to isolate sources of mechanical properties can be confounding since protein complexes that generate forces, e.g., actomyosin can be the same as the ones transmitting force and establish tissue elasticity (Zhou et al., 2009; Zhou et al., 2010; Zhou et al., 2015). These overlapping roles make it challenging to isolate elasticity from force generation. Furthermore, the stochastic nature of force generation can obscure cells’ passive deformation and hide the true sources of heterogeneity. In this project we make use of the *Xenopus* embryonic ectoderm, a tissue that does not generate a tissue-scale morphogenetic force. Thus, stretching Xenopus embryonic ectoderm provides us with a passive tissue where we can externally drive "morphogenesis" with a tissue stretcher and modulate cell biology that is involved in morphogenesis but without directed forces that active tissues would generate.

In this study, we used a recently published tissue stretcher to apply high uniaxial mechanical strains up to 100% on organotypic epithelial explants from developing *Xenopus laevis* embryos to test how mechanical forces are transmitted within the cells (Yang et al., 2025). Using a combination of techniques including live imaging, tension-sensors, laser ablation, and perturbations of actomyosin networks, we identify how mechanical forces are transmitted through epithelial cells. We show that epithelial cells respond heterogeneously to applied force, with cell-to-cell variation and test contributors to these responses. We find Poisson’s ratio emerges as a key determinant of strain heterogeneity, suggesting that epithelial cells behave as an anisotropic, rather than homogeneous material. Through vinculin force sensing and laser ablation, we implicate the medio-apical actomyosin cortex as the primary force transmitter, with junctional actomyosin playing a role in force dissipation and remodeling. Together, these findings reveal how epithelial tissues adapt to mechanical forces by integrating junctional and cortical actomyosin networks.

## 2. Results

### 2.1. Cells within the tissue respond heterogeneously to externally applied tension

To visualize cell response to externally applied tension, we use a previously described customized tissue stretcher (TissueTractor; (Yang et al., 2025)) to apply defined uniaxial strain on *Xenopus laevis* epithelial organotypic explants (e.g., animal cap). In brief, the TissueTractor consists of two linear actuators, a 3D-printed microscope stage insert compatible with inverted microscopes, and a cassette for tissue attachment (**Figure 1a, and 1a’**). Explants attach to the underside of the elastic polydimethylsiloxane (PDMS) substrate on the cassette (**Figure 1a’**). The explants were first imaged in a relaxed state (S_0_) and incrementally stretched, with images captured at each step (S_1_, S_2_, …, S_n_). The TissueTractor allows tracking the same group of cells over the entire course of tissue stretching (**Figure 1b**).

**Figure 1.**
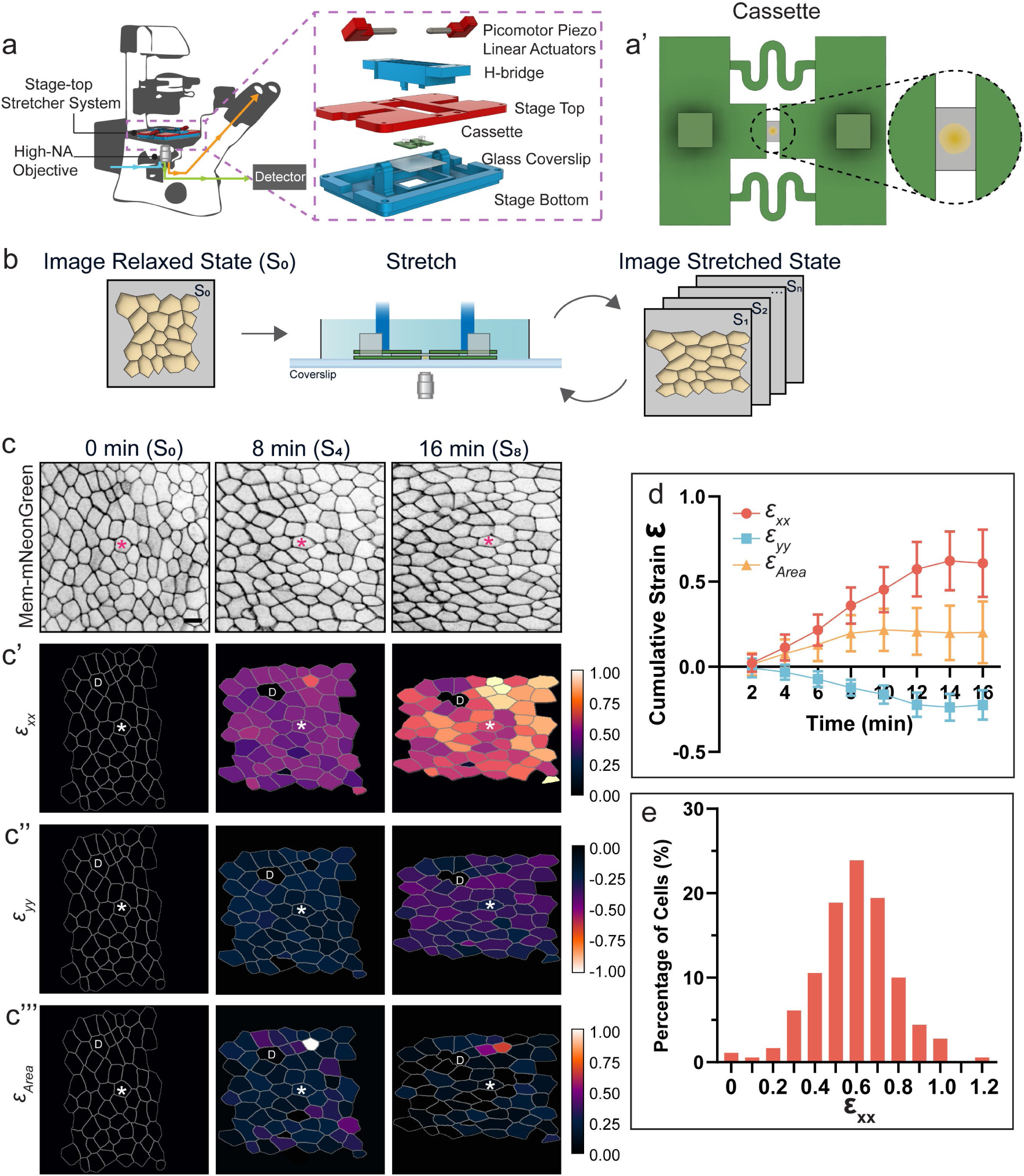
Tissue stretcher set up and quantitation of strain heterogeneity under externally applied tension. (**a**) A schematic of the stage-top stretcher system TissueTractor on an inverted microscope. The dashed box shows an exploded view of all components in the TissueTractor. From top to bottom: two picomotor piezo linear actuators, and “H-bridge” with extended cantilevers controlled by the movement of actuators, a stage top, a disposable cassette, and a stage bottom including a 45 x 50 mm glass coverslip. (**a’**) A schematic of the disposable cassette. The dashed circle zooms in on an organotypic explant attached on the underside of the PDMS substrate that spans in between the PES shims. (**b**) The workflow of imaging and stretching experimental setup. (**c**) Representative images of a Stage 13 – 14 animal cap organotypic explant labeled with membrane-mNeonGreen at 0 min (S_0_, relaxed state), 8 min (S_4_), and 16 min (S_8_). * indicates the same cell across frames. Scale bar = 20 μm. (**c’**) ɛ_xx_ strain (**c’’**) ɛ_yy_ strain, and (**c’’’**) ɛ_Area_ strain mapped using individual cellular strain at 0 min, 8 min, and 16 min. At 0 min, cells were at relaxed state, which all cells have strain of 0 (shown in black with grey cell outlines). Calibration bars for ɛ_xx_ and ɛ_Area_ strain ranges from 0 to 1, with 0 in black and 1 in white. Calibration bar for ɛ_yy_ strain ranges from -1 to 0, with 0 in black and -1 in white. * indicates the same cell across frames. “D” indicates the cell divided during stretching, which was not included in strain calculation. N = 60 cells. (**d**) Quantifications of cellular strain ɛ_xx_, ɛ_yy_, and ɛ_Area_ at each stretch step (2 min, 4 min, …, 16 min). Error bars, standard deviation. N = 181 cells. (**e**) Frequency distribution of cells in binned ɛ_xx_ strain at 16 min. Individual bin size is 0.1. N = 181 cells. (**a**) and (**a’**) are adapted from Yang et al. 2025 (Yang et al., 2025).

To start, we stretched an epithelial explant in 8 steps (S_0_ to S_8_), and imaged at 2-minute interval between consecutive increments. By the end of stretch, the grip-to-grip engineering strain (**Box 1**) applied in the cassette reached 1.5, corresponding to an average strain rate of 0.094 per minute (Figure 1c) (Yang et al., 2025). In this case, tissue in the explant reached a maximum strain of 0.66 ± 0.047 along the stretch axis (x-axis; **Figure S1**), resulting in a tissue strain rate (**Box 1**) of 0.04 ± 0.003 per minute. To quantify individual cellular strain, cells were segmented (Mashburn et al., 2012) and engineering strain in each cell was calculated including ɛ_xx_ (along the stretch axis; x-axis), ɛ_yy_ (perpendicular to the stretch axis; y-axis), and area strain ɛ_Area_ (Vijayraghavan, 2018; Yang et al., 2025). During the stretch, individual cellular strains ɛ_xx_ became increasingly heterogeneous during stretching, reaching 0.61 ± 0.20 at the end of the experiment (16 min), resulting in a mean cell strain rate of 0.04 ± 0.012 per minute along the x-axis (**Figure 1c’ and 1d**). As expected under uniaxial strain, cells compress along y-axis (ɛ_yy_ = - 0.22 ± 0.087) (**Figure 1c’’ and 1d**). Surprisingly, the area strain ɛ_Area_ was also positive (ɛ_Area_ = 0.20 ± 0.18), indicating cell apical areas increased an average of 20% despite compression along the y-axis (**Figure 1c’’’, and 1d**). Such an anisotropic deformation may be due to the compression in the apical-basal direction (e.g., z-axis) exceeding the compression along the y-axis. If we assume cell volume is constant, we can infer that cells are stiffer in the epithelial plane than along the apical-basal axis.

#### Box 1. Glossary

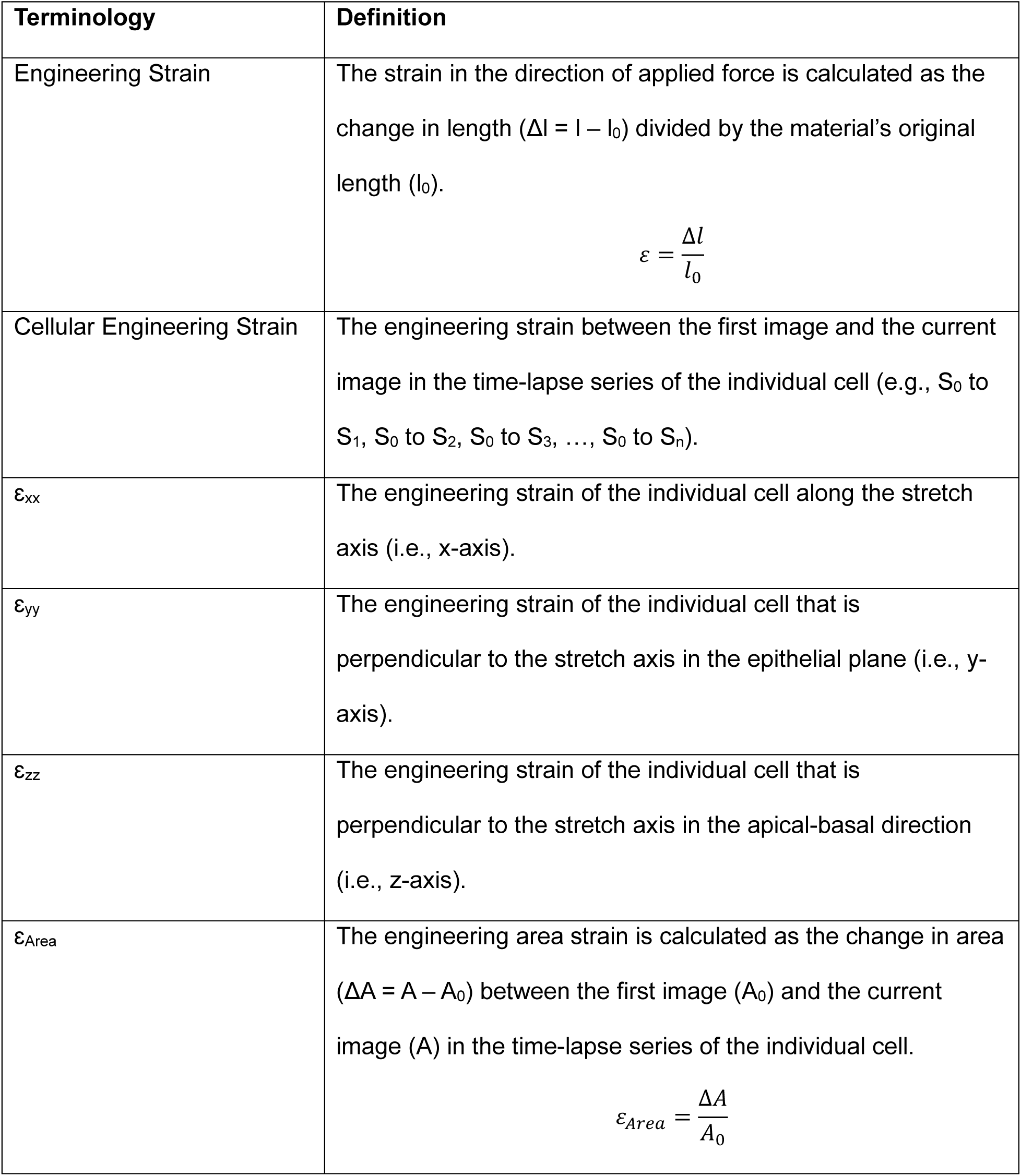

| Terminology | Definition |
| --- | --- |
| Engineering Strain | The strain in the direction of applied force is calculated as the change in length ($\Delta l = l - l_0$) divided by the material's original length ($l_0$).<br>$\varepsilon = \frac{\Delta l}{l_0}$ |
| Cellular Engineering Strain | The engineering strain between the first image and the current image in the time-lapse series of the individual cell (e.g., $S_0$ to $S_1$, $S_0$ to $S_2$, $S_0$ to $S_3$, ..., $S_0$ to $S_n$). |
| $\varepsilon_{xx}$ | The engineering strain of the individual cell along the stretch axis (i.e., x-axis). |
| $\varepsilon_{yy}$ | The engineering strain of the individual cell that is perpendicular to the stretch axis in the epithelial plane (i.e., y-axis). |
| $\varepsilon_{zz}$ | The engineering strain of the individual cell that is perpendicular to the stretch axis in the apical-basal direction (i.e., z-axis). |
| $\varepsilon_{Area}$ | The engineering area strain is calculated as the change in area ($\Delta A = A - A_0$) between the first image ($A_0$) and the current image ($A$) in the time-lapse series of the individual cell.<br>$\varepsilon_{Area} = \frac{\Delta A}{A_0}$ |

Since ɛ_xx_ exhibited greater heterogeneity (standard deviation = ± 0.20) than ɛ_yy_ (standard deviation = ± 0.087), we focused on distribution of ɛ_xx_ strain. We note that strain heterogeneity is not due to the variation in strain of the PDMS elastic substrate; our previous study confirmed a high degree of strain uniformity in PDMS substrates used in the TissueTractor cassettes (Yang et al., 2025). Thus, applied strain is uniform across the substrate. Therefore, the observed variability in ɛ_xx_ arises from variations in intrinsic cellular properties. At the end of the series of stretches (16 min), the frequency distribution of ɛ_xx_ showed that 62% of cells reached 0.50 to 0.70 engineering strain, with 20% exhibiting lower strain and 18% exceeding this range (**Figure 1e**). Additionally, strain heterogeneity did not resolve as tissues were held in the stretched configuration. After one hour, we found that ɛ_xx_ of the cells slightly relaxed along x- and y-axes, with ɛ_xx_ decreasing from 0.64 ± 0.19 at 16 min to 0.53 ± 0.19, and ɛ_yy_ also relaxed from −0.26 ± 0.10 to −0.19 ± 0.15, while ɛ_Area_ was unchanged (**Figure S2a**). However, even as cells relaxed (e.g., ɛ_xx_ strain shifted 0.10 lower) they remained heterogeneous (**Figure S2b, and S2c**). These findings indicate that epithelial tissues behave as a heterogeneous material, rather than a homogeneous one, with individual cells responding variably but consistently to the same applied mechanical strain.

It has been suggested that variation in epithelial responses to strain can arise from multiple factors, including differences in cell geometry, mechanical properties, ECM compositions, cytoskeletal organization, and force transmission within the tissue (Gibson et al., 2006; Munjal and Lecuit, 2014; Oakes et al., 2014; Bachir et al., 2017; Chanet et al., 2017; Bodenschatz et al., 2022). However, these studies have not directly tested the specific contributions of these factors to strain heterogeneity.

### 2.2. Cell geometry contributes to cellular strain heterogeneity

To test the sources of strain heterogeneity within epithelial tissues, we first investigated the roles of cell geometry and intrinsic material properties in shaping cellular strain responses. Since previous studies suggest that both cell geometry and material properties influence cellular mechanics (Gibson et al., 2006; Oakes et al., 2014; Bi et al., 2016; Li et al., 2016; Chanet et al., 2017; Bodenschatz et al., 2022), we hypothesized that initial cell geometry might be correlated to the emergence of ɛ_xx_ heterogeneity (**Figure 2a**). To test this, we examined the initial conditions of cell apical surface area, cell shape index, aspect ratio, cell orientation, and coordination number, and assessed their correlations with ɛ_xx_ strain.

**Figure 2.**
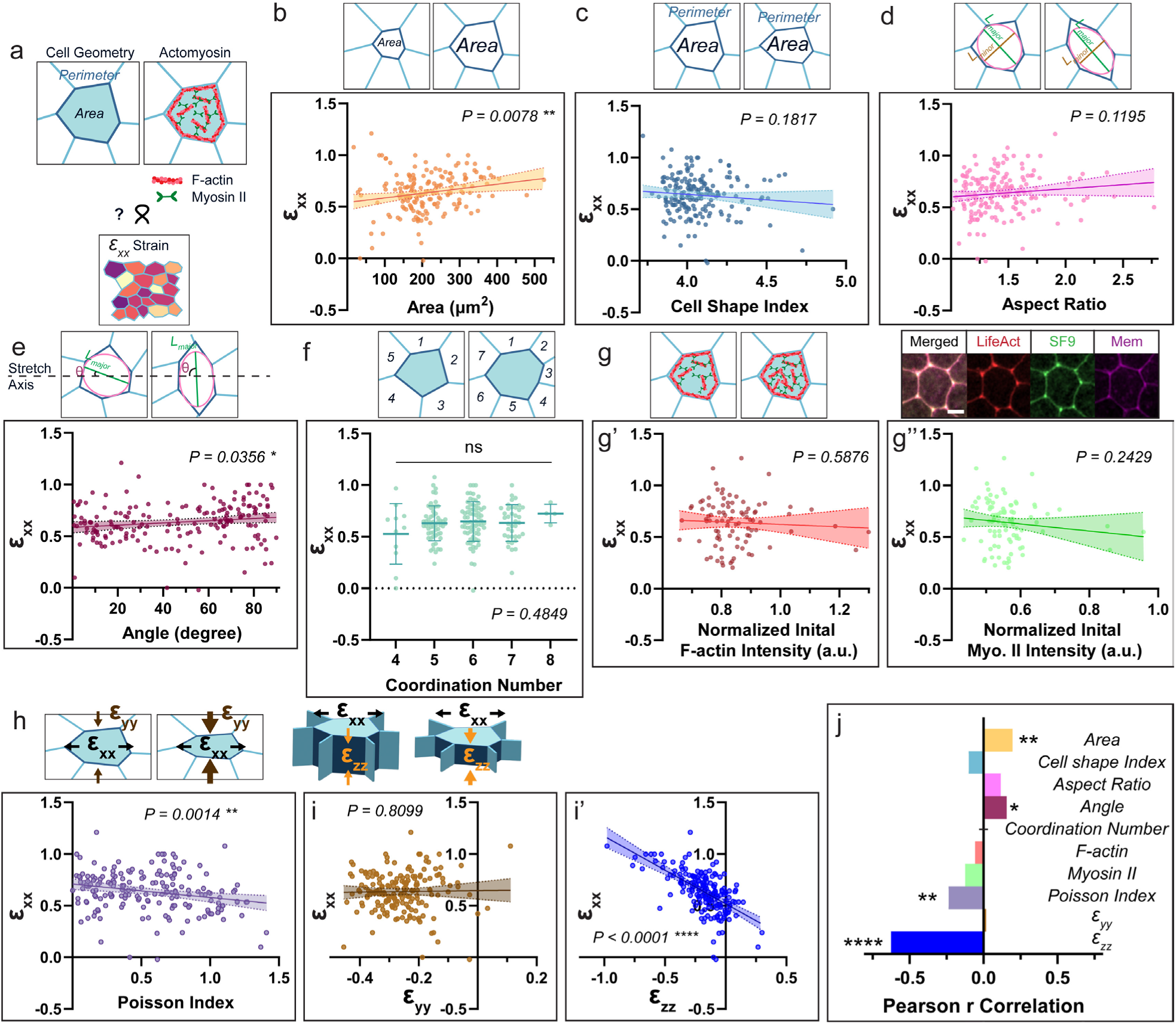
Geometric contributors to strain heterogeneity under externally applied tension. (**a**) The schematic shows that cell geometry and actomyosin levels may correlate with ɛ_xx_ strain heterogeneity. (**b**) Cell surface area shows significant positive correlation to ɛ_xx_ strain. (**c**) Cell shape index, (**d**) aspect ratio, calculated as the ratio between length of the major axis (green) and length of the minor axis (brown) of the largest fitted ellipse (outlined in pink) in the cell, and (**e**) The orientation of the cell shows significant positive correlation to ɛ_xx_ strain. (**f**) Coordination numbers (Error bars, standard deviation) show no significant correlation to ɛ_xx_ strain. (**g**) The schematic shows cells with actomyosin components. A representative cell fluorescently labeled with an F-actin marker LifeAct, a Myosin II marker SF9, and a membrane marker. Scale bar = 10 μm. (**g’**) Normalized LifeAct intensity, and (**g’’**) normalized SF9 intensity in the individual cells at relaxed state shows no significant correlation to ɛ_xx_. N = 92 cells. (**h**) The schematic shows that a cell experiences compressive strain along yy direction (ɛ_yy_, brown) and zz direction (ɛ_zz_,orange) when stretched in xx direction (ɛ_xx_). Poisson Index shows significant negative correlation to ɛ_xx_. (**i**) ɛ_yy_ shows no correlation to ɛ_xx_. (**i’**) ɛ_zz_ shows significantly negative correlation to ɛ_xx_. N = 181 cells for panels (**b-f, h, and i**). (**j**) Summary plot of Pearson correlation coefficient r calculated from correlation between each parameter and ɛ_xx_. Positive values indicate positive correlation whereas negative values indicate negative correlation to ɛ_xx_. The length of the bar indicates the degree of correlation. Correlation between each parameter and ɛ_xx_ is calculated using simple linear regression except the number of cell neighbors. Linear regression lines are in solid colors, and the dashed lines with shades in between represent the 95% confidence bands. Schematics of each parameter are shown in **b - i**, with the right schematic representing a larger value of the specific parameter compared to the left schematic.

Apical surface area shows a positive correlation with ɛ_xx_, where cells with initially larger surface area exhibiting greater strains than cells with smaller surface area (**Figure 2b, and 2j**). Next, we tested whether cell shape parameters correlate to strain heterogeneity, as prior research suggested that cells with smaller cell shape index exhibit are more solid-like (Bi et al., 2016; Erdemci-Tandogan et al., 2018; Jain et al., 2020). Cell shape index is calculated as:

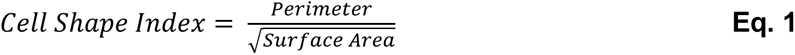

where perimeter is the length of the path outlines the cell, and the surface area is the measure of the size of the apical surface of the cell. However, we found no significant correlation between the initial cell shape index and ɛ_xx_ (**Figure 1c**). Another common cell shape parameter is the aspect ratio:

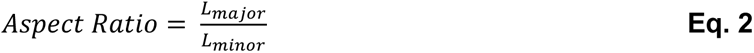

where L_major_ is the length of the major axis (long axis) of the ellipse, and L_minor_ is the length of the minor axis (short axis) of an ellipse fitted to a segmented cell outline (**Figure 2d**). Again, we observed no significant correlation with ɛ_xx_ (**Figure 2d, and 2j**). We then tested the cell orientation, as prior work indicated that stress propagation is more efficient when the cells are perpendicular to the direction of stress (Ruppel et al., 2023). We found a small positive correlation between the initial cell orientation and ɛ_xx_, indicating that cells oriented perpendicular to the direction of strain experienced greater strain (**Figure 2e, and 2j**). These results suggest that while cell surface area and orientation influence strain, shape parameters alone do not strongly predict strain heterogeneity.

We then considered coordination number, defined as the number of neighbors that each cell contacts (O’Keeffe, 1979). Strain distribution may depend on local cell connectivity, with cells with a coordination number of 6 being more mechanically stable in metazoan epithelia (Gibson et al., 2006). We hypothesized that cells with more neighbors may experience lower strain since forces redistribute among more neighboring cells. Most cells in *Xenopus* ectoderm here have 5 to 7 neighbors; we found no significant correlations between number of neighbors and ɛ_xx_ (**Figure 2f**).

### 2.3. Pre-strain initial actomyosin levels do not contribute to strain heterogeneity

Cells within the same tissue vary in initial actomyosin levels, and it is commonly assumed that cells with higher actomyosin levels are stiffer (Kollimada et al., 2021). We hypothesized that cells with higher initial actomyosin levels would resist deformation and exhibit lower strain. To test this, we co-expressed fluorescently-tagged LifeAct and SF9 intrabody (Nizak et al., 2003), to label F-actin, and active myosin II, respectively, together with a cell membrane marker (**Figure 2g, and 2j**). Neither initial F-actin nor active myosin levels in the cell correlated with ɛ_xx_ (**Figure 2g’, 2g’’, and 2j**), suggesting that pre-existing actomyosin levels do not predict cellular strain responses under external tension.

### 2.4. Aniostropic structural properties of the cell contribute to strain heterogeneity

Given our earlier findings of anisotropic cellular responses along different axes (**Figure 1c, and 1d**), we next investigated the Poisson effect. From material science, Poisson effect is determined by material internal structure, such as atomic packing and alignment (Greaves et al., 2011). The Poisson effect describes the deformation in material in a direction perpendicular to the direction of the applied force. In isotropic, incompressible materials, stretching along the x-axis results in uniform compressions in y- and z-directions. In anisotropic materials, Poisson effect depends on material orientation (Greaves et al., 2011). We have shown that epithelial sheets exhibit anisotropy along y- and z-axes under uniaxial tension along x-axis (**Figure 1c, and 1d**). To investigate the Poisson effect, we calculated Poisson’s ratio *ν*_xy_ in the xy plane (i.e., the epithelial plane), and *ν*_xz_ in the xz plane (i.e., the apical-basal plane):

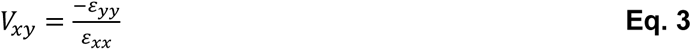

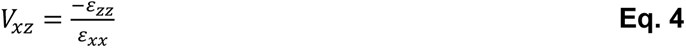

where ɛ_xx_, ɛ_yy_, and ɛ_zz_ are defined in **Box 1**. We found a Poisson’s ratio of *ν*_xy_ = 0.43 ± 0.19 and *ν*_xz_ = 0.26 ± 0.22, indicating the apical-basal (xz plane) direction is relatively stiffer than the epithelial (xy plane) plane. One possible explanation is that structural elements aligned within the epithelial plane are more easily compressed than those along the apical-basal axis. To further assess whether this anisotropy contributes to strain heterogeneity, we created Poisson Index, which is an anisotropic index normalized by their Euclidean norm, independent of ɛ_xx_:

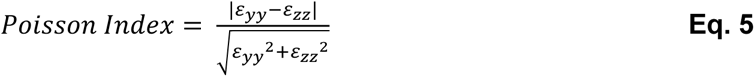

In an isotropic, incompressible material, the Poisson Index should equal 0. We found a significant negative correlation between Poisson Index and ɛ_xx-_ (**Figure 2h, and 2j**), suggesting cells with greater anisotropy experience more strain along the x-axis. Next, we investigated whether the strain along y-axis and z-axis contribute to strain heterogeneity along x-axis. While ɛ_yy_ has no significant correlation to ɛ_xx_ (**Figure 2i, and 2j**), ɛ_zz_ shows a strong negative correlation to ɛ_xx_ (**Figure 2i’, and 2j),** indicating that greater compression in the apical-basal plane corresponds to greater strain along the x-axis. These findings suggest that structural properties of the cell contribute to strain heterogeneity, with epithelial anisotropy playing a key role. Unlike isotropic materials, where mechanical responses are uniform in all directions, epithelial tissues exhibit direction-dependent mechanical behavior, likely due to differences in polarity along different axes that directs cytoskeletal assembly and establishes the architecture of cell-cell and cell-substrate junctions.

### 2.5. Actomyosin remodeling during stretch reflects distinct roles in force transmission and dissipation

While we demonstrated that initial, pre-strain actomyosin levels do not dictate cell strain heterogeneity (**Figure 2i**), actomyosin remodeling is thought to play a critical role in cellular responses to mechanical stress (Sumi et al., 2018; Khalilgharibi et al., 2019). To investigate actomyosin changes under mechanical strain, we stretched tissues expressing F-actin, myosin II, and membrane markers (**Figure 3a**), and measured actomyosin intensities before and after stretch. Both F-actin and myosin II intensities increased after stretch (**Figure 3b**), consistent with observations in basal actomyosin in follicle cell expansion and in ectoderm during convergent extension in *Drosophila* embryo (Kale et al., 2018; Li et al., 2024). We further investigated whether this increase in actomyosin correlated with the magnitude of individual cell strain and found positive correlation between ɛ_xx_ strain and changes in actomyosin (**Figure 3c, and 3d**). Thus, cells that experience greater strain show larger increases in actomyosin levels, suggesting that actomyosin is not the cause of strain heterogeneity but rather a graded response to mechanical strain.

**Figure 3.**
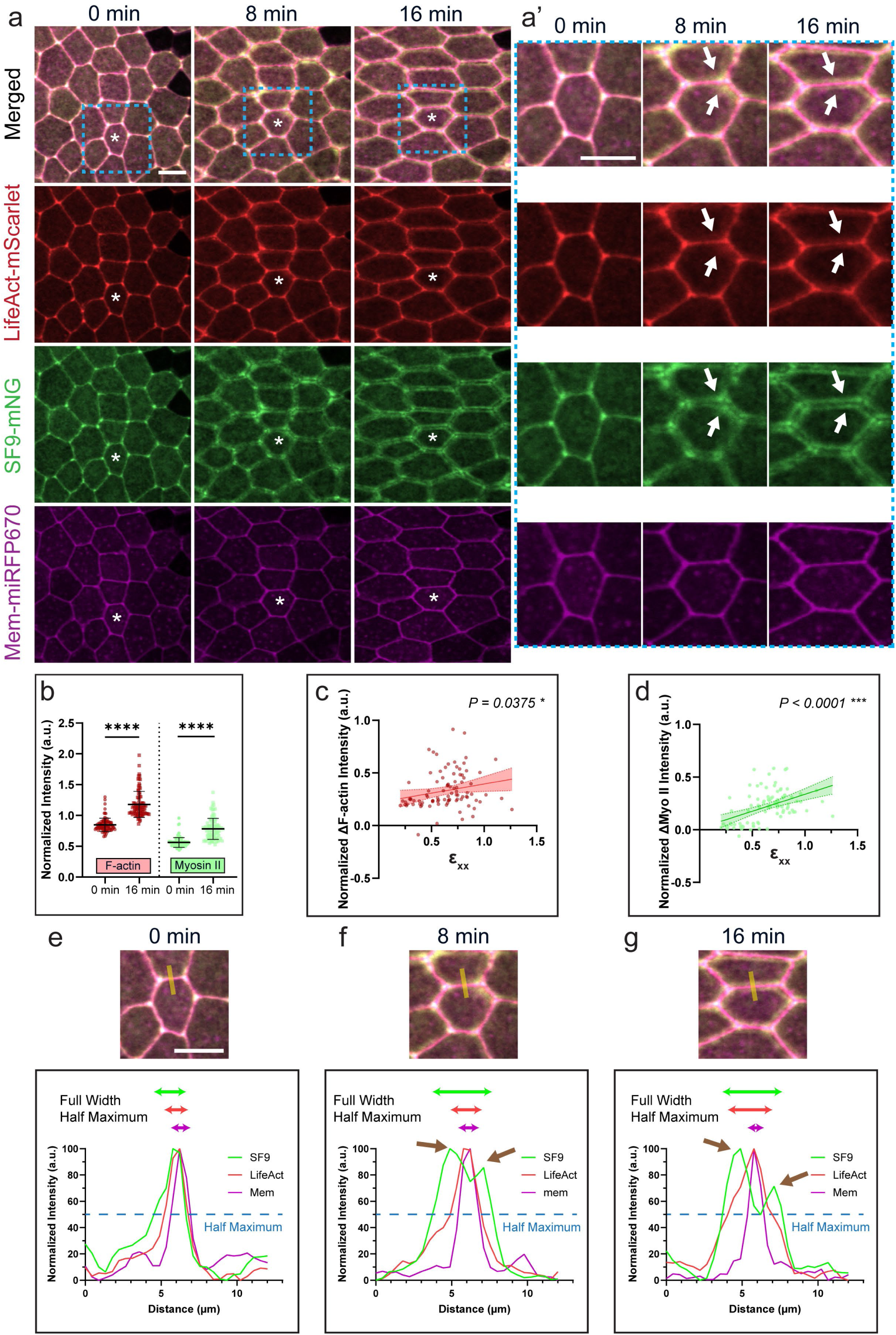
Applied strain induces spatial reorganization of actomyosin. (**a**) Representative images of *Xenopus* epithelial tissues expressing LifeAct-mScarlet, SF9-mNeonGreen, and membrane-miRFP670 at 0 min (relaxed), 8 min, and 16 min. * indicates the same cell throughout frames. Cyan dashed box indicates the zoom-in region in (**a’**). White arrows show the F-actin flare and myosin II bands at stretched state (8 and 16 min). (**b**) Quantifications of F-actin and myosin II intensity at relaxed (0 min) and stretched (16 min) states, with both significantly increased after stretch. (**c**) F-actin, and (**d**) myosin II both positively correlated with strain ɛ_xx_. (**e**) Intensity profiles of membrane, LifeAct, and SF9 across a cell junction at 0 min, (**f**) at 8 min, (**g**) and at 16 min. Yellow lines show the location of plotted intensity profiles across the junction. Brown arrows indicate the two separate intensity peaks of myosin II at 8 and 16 min. The blue dashed line represents half of the maximum intensity. Green, red, and magenta horizontal lines at the top of the plot represent the width of each intensity band at its half maximum (Full Width Half Maximum). Scale bar = 20 μm.

Upon stretching, myosin II accumulated in the medio-apical surface, forming distinct bands that deviated from cell junctions, leaving a gap at the junctions (**Figure 3a’, white arrows**). F-actin remained prominent at cell junctions, diffused F-actin signals were also observed in the medio-apical cortex (**Figure 3a’, white arrows**). At the relaxed state, intensity profiles across the junction showed single, narrow peaks for membrane, F-actin, and myosin II channels (**Figure 3e**). However, under stretched conditions (8 min and 16 min), myosin II profiles displayed two distinct peaks flanking the membrane and F-actin peaks (**Figure 3e, and 3f, brown arrows**), indicating a shift of myosin II away from the cell junction. Although F-actin retained a single peak centered at the junction, its localization band broadened after stretch, as shown by an increase in full width at half maximum (FWHM), whereas membrane signals remained sharply confined during stretching (**Figure 3e – g**). Similar banding in actomyosin structures have been reported during wound healing in *Xenopus* oocytes, *Drosophila* eye morphogenesis, and after β-catenin RNAi-mediated silencing in *Drosophila* embryos (Benink and Bement, 2005; Curran et al., 2017; Rosa-Birriel et al., 2024). However, none of these studies have associated this medio-apical actomyosin accumulation directly with mechanically induced strain. Given this distinct actomyosin reorganization under tension, we further investigated the dynamics of junctional and medio-apical actomyosin.

To investigate the dynamics of junctional and medio-apical actomyosin we performed fluorescence recovery after photobleaching (FRAP) on junctional and medio-apical LifeAct that labeled F-actin (Arnold et al., 2019) in both relaxed and stretched tissue (**Figure 4a**). FRAP measures three key biophysical properties: mobile and immobile fractions, representing the portion of actomyosin that recovers and does not recover after bleaching, respectively, and the half-life, defined as the time required to recover half of the maximum recovered fluorescence intensity (Sprague and McNally, 2005). We found that in both relaxed and stretched tissues, the medio-apical F-actin recovered significantly less than the junctional F-actin (**Figure 4b**), with a consistently higher immobile fraction (**Figure 4d, and 4e**), suggesting medio-apical F-actin is inherently more stable than junctional F-actin. Furthermore, upon stretching, both junctional and medio-apical F-actin became more mobile (**Figure 4b, 4d, and 4e**) suggesting that mechanical tension destabilizes F-actin across the cell, in both medio-apical and junctional regions.

**Figure 4.**
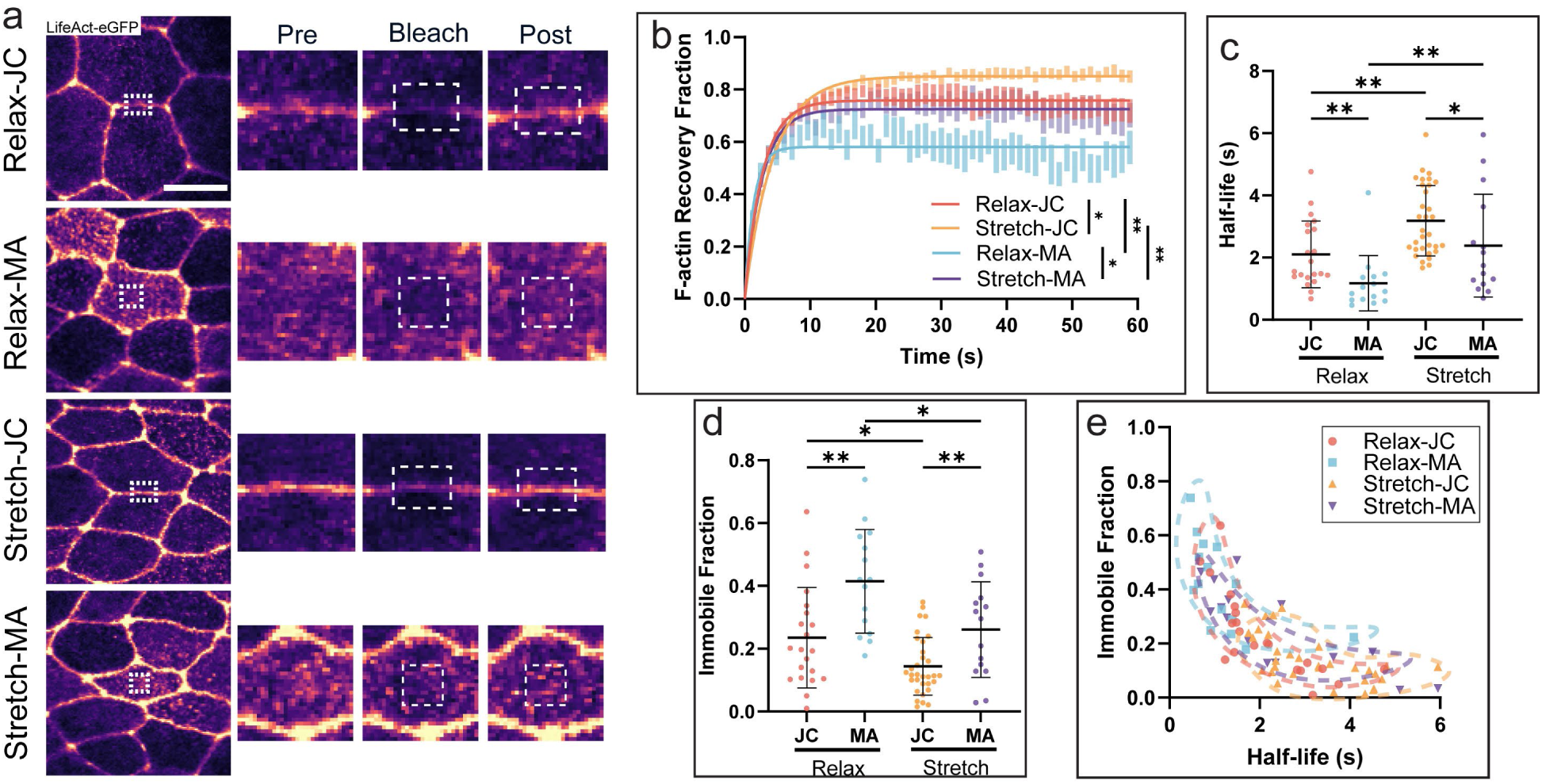
Applied strain destabilizes both junctional and medio-apical F-actin networks, with medio-apical F-actin exhibiting greater stability. (**a**) Representative images of FRAP on junctions and medio-apical surfaces in relaxed (Relax-JC (N = 21), Relax-MA (N = 14)) and stretched (Stretch-JC (N = 32), Stretch-MA (N = 15)) tissues. Zoom-in regions around the FRAP regions at pre-bleach, bleach, and post-bleach are shown on the right. White dashed boxes indicate the bleached regions. (**b**) Normalized F-actin recovery curve after photobleaching. (**c**) Quantification of half-life, and (**d**) immobile fraction in FRAP for JC and MA at relaxed and stretched states. (**e**) Half-life vs. immobile fraction of JC and MA at relaxed and stretched states. Scale bar = 20 μm.

FRAP recovery curves further reveal that medio-apical F-actin exhibited a significantly shorter half-life than junctional F-actin in both relaxed and stretched conditions (**Figure 4c, and 4e**), indicating that the fraction of actomyosin that does recover in the medio-apical domain is remodeled more quickly. This suggests that even though apical cortex structures are stable, there is a more dynamic turnover rate of actomyosin in the medio-apical surface. Comparing relaxed and stretched conditions, the half-life of junctional F-actin increased under stretch, whereas medio-apical F-actin did not (**Figure 4c, and 4e**).

Together, these findings indicate that medio-apical and junctional actomyosin respond differently to mechanical strain, with medio-apical actomyosin remaining more stable (**Figure 4e**). This difference in stability and mobility suggests distinct functional roles, with medio-apical actomyosin likely playing a key role in force transmission, as junctional actomyosin facilitating force dissipation.

### 2.6. Medio-apical actomyosin as a key driver of force transmission during stretch

To further explore whether forces are transmitted via medio-apical domain of cells or cell-cell junction complexes, we examined vinculin recruitment as a readout for relative tension. In epithelial cells, vinculin can function as a mechanosensor (Sala et al., 2024); under tension, F-actin bound to α-catenin at cell-cell junctions induces a conformational change that exposes a binding site for vinculin (**Figure 5a, and 5a’**). Thus, vinculin is recruited to sites experiencing tension (Collinet and Lecuit, 2013; Arnold et al., 2019; Chirasani et al., 2023). Since junctional and medio-apical actomyosin can both act as a source of tension applied on α-catenin to recruit vinculin (**Figure 5a, and 5a’**), we hypothesized that if junctional actomyosin is the primary source of force transmission during stretch, vinculin would be recruited more to the junctions parallel to the stretch axis (**Figure 5a**). Alternatively, if medio-apical actomyosin is the primary source, vinculin would be recruited more to the junctions perpendicular to the stretch axis (**Figure 5a’**). To test this, we stretched epithelial tissues expressing vinculin and membrane markers (**Figure 5b**).

**Figure 5.**
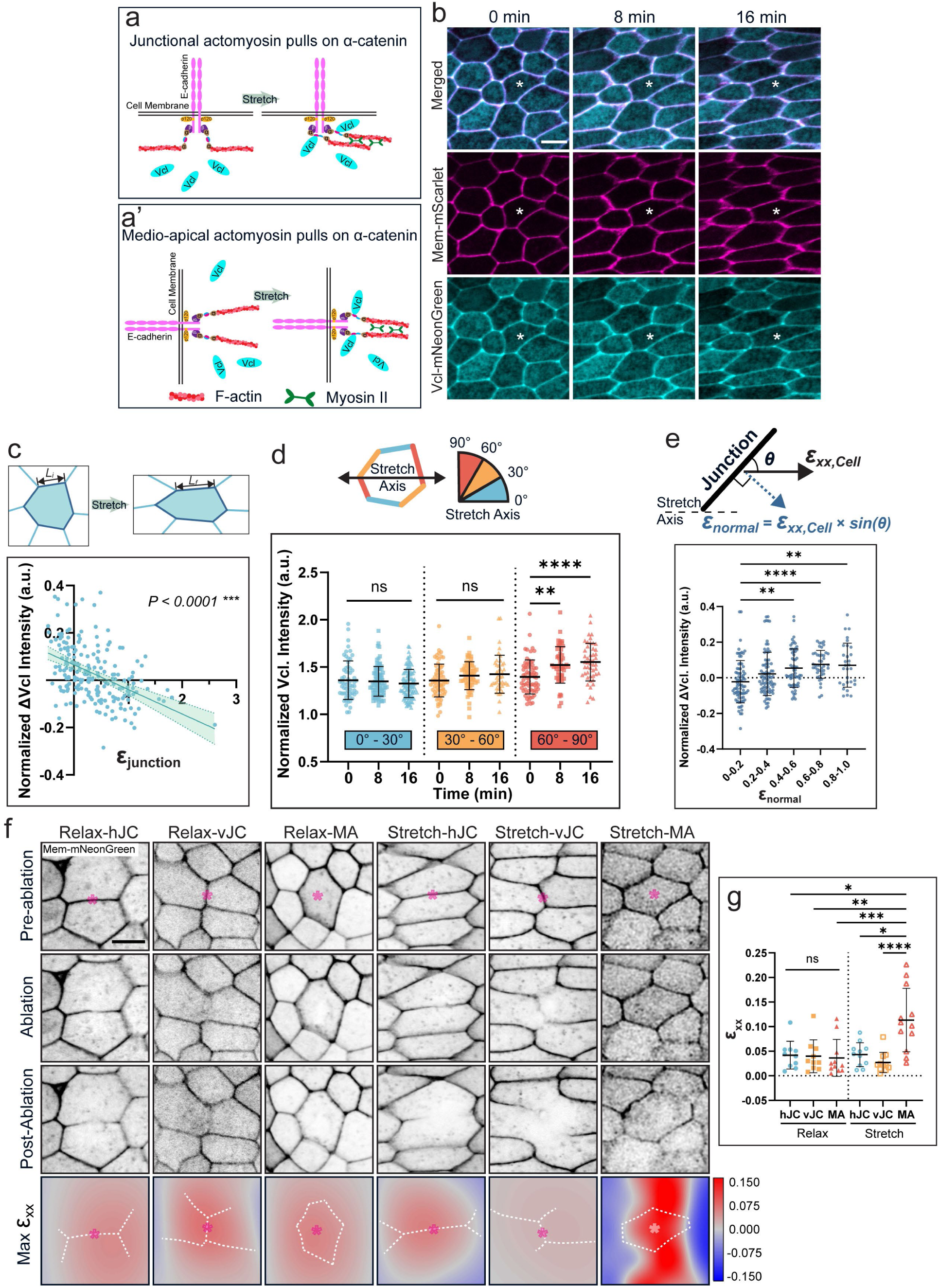
Medio-apical cortex is the major mechanical structure of the apical epithelium: vinculin recruitment and laser ablation. (**a**) A schematic shows vinculin recruitment to α-catenin vinculin binding sites upon stretch via junctional actomyosin. (**a’**) A schematic shows medio-apical actomyosin pulls on α-catenin to recruit vinculin to the junction. Double black lines: cell membrane; pink: E-cadherin; yellow: p120 proteins; purple: β-catenin; brown with blue and magenta: α-catenin; Cyan: vinculin. (**b**) Representative images of *Xenopus* epithelial tissues expressing vinculin-mNeonGreen and membrane-mScarlet at 0 min (relaxed), 8 min, and 16 min. * indicates the same cell throughout frames. (**c**) A schematic shows junction length increases after stretch. ɛ_junction_ has a negative correlation to the normalized vinculin intensity changes. N = 199 junctions. (**d**) A schematic represents cell junctions at three angle categories respective to the stretch axis: 0° - 30° (blue), 30° - 60° (yellow), and 60° - 90° (red). Quantifications of normalized vinculin intensity from three groups at 0 min (relaxed), 8 min, and 16 min). N = 199 junctions. (**e**) A schematic shows that the normal component of the cell strain ɛ_normal_ is calculated by the cell strain ɛ_xx,Cell_ multiplies the sine of the angle between the junction and the stretch axis. Quantifications of normalized vinculin intensity changes at increasing ɛ_normal_ strains. (**f**) Representative images of laser ablations performed on parallel junctions (hJC), perpendicular junctions (vJC), and medio-apical (MA) regions in relaxed and stretched tissues. Respective maximum ɛ_xx_ strain maps of the ablated sites are shown in the last row. * indicates the ablation site. White dashed lines represent the ablated junctions or cells. (**g**) Quantifications of maximum ɛ_xx_ at the ablation site of hJC, vJC, and MA in relaxed and stretched tissues. N = 10 (Relax-hJC), 10 (Relax-vJC), 11 (Relax-MA), 11 (Stretch-hJC), 11 (Stretch-vJC), 12 (Stretch-MA). Scale bar = 20 μm.

We first examined whether vinculin recruitment is affected by junctional strain ɛ_junction_:

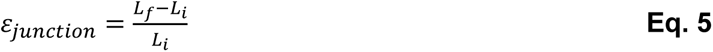

where L_i_ is the initial length of the junction before stretch, and L_f_ is the finial length of the junction after stretch. We found a strong negative correlation between vinculin intensity changes and junctional strain, indicating that decreasing junctional strain led to increased vinculin recruitment (**Figure 5c**). This finding is particularly intriguing, as junctions parallel to the stretch axis are expected to experience the greatest tensile forces and length changes under uniaxial strain. However, our results indicate that vinculin was not preferentially recruited to these high-strain junctions, suggesting that force transmission occurs through an alternative mechanism.

Next, we grouped all cell junctions into three angle categories relative to the stretch axis: 0° to 30° (parallel, blue), 30° to 60° (intermediate, yellow), and 60° to 90° (perpendicular, red) (**Figure 5d**). We then measured normalized vinculin intensity along these junctions during stretch. We found a significant increase of vinculin intensity along only the perpendicular junctions during stretch (**Figure 5d**). In addition, at 8 min, and 16 min stretched steps, vinculin intensity along junctions perpendicular to the stretch axis was higher than along junctions parallel to the stretch axis (**Figure S3)**.

Thus, medio-apical tension, rather than junctional tension, recruits vinculin tension sensors during uniaxial stretch. We calculated the normal strain component ɛ_normal_ from the individual cellular strain ɛ_xx,Cell_ :

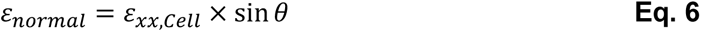

where θ is the angle between the junction and the stretch axis (**Figure 5e**). We found that as ɛ_normal_ increased, vinculin localization increased, plateauing after ɛ_normal_ reached 0.6 (**Figure 5e**). This plateau may result from most α-catenin along the junctions reaching full conformational activation at 0.6 strain, saturating vinculin recruitment as all available α-catenin sites become occupied.

To conceptualize these findings, we developed an idealized model of the apical cortex of an epithelial cell. The ideal cell is represented as a square unit containing only two types of junctions: those parallel to the stretch axis and those perpendicular to the stretch axis. In this model, actomyosin behaves as a series of mechanical springs, where blue springs correspond to junctional actomyosin and red springs represent medio-apical actomyosin (**Figure 6b**). This model aligns with our earlier hypothesis (**Figure 5a and 5a’**), where force transmission is either dominated by junctional actomyosin (**Figure 5a**) or medio-apical actomyosin (**Figure 5a’**). Under uniaxial tension, we observed no enrichment along junctions parallel to the stretch axis, ruling out junctional actomyosin as the primary force transmitter (**Figure 5d**). Vinculin enriches exclusively along the perpendicular junctions (**Figure 5d**), supporting the conclusion that force transmission primarily occurs through medio-apical actomyosin rather than junctional actomyosin.

**Figure 6.**
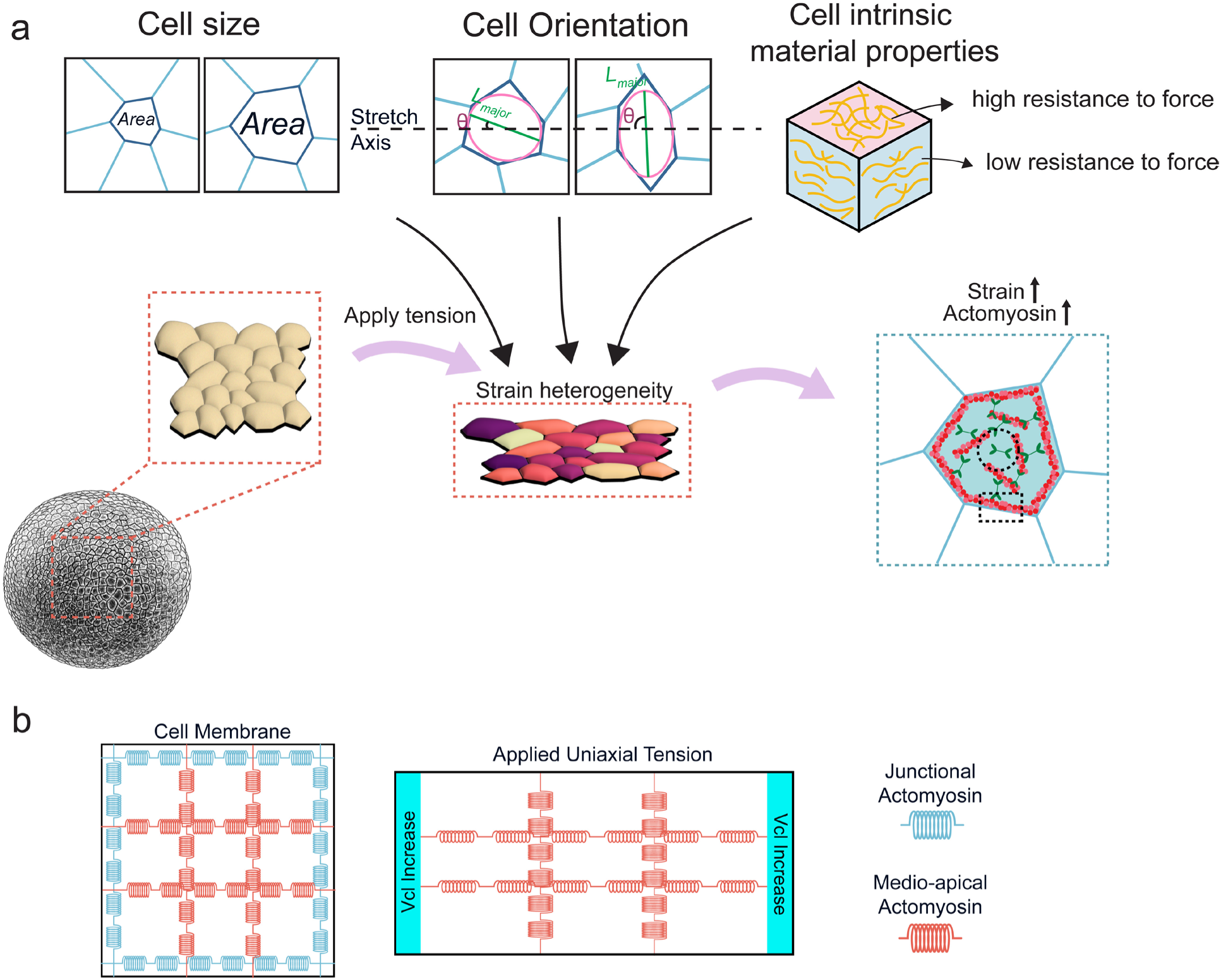
A schematic representation of heterogeneous strain responses in epithelial tissues where responses are governed by cell geometry, intrinsic material properties and medio-apical actomyosin dynamics. (**a**) Upon applied tension, cells within the same tissue exhibit variable strains, influenced by cell surface area, cell orientation, and cell intrinsic material properties. This heterogeneous strain responses lead to differential actomyosin upregulation, where cells experiencing higher strain exhibit greater increases in actomyosin levels. (**b**) A model of a square unit cell with actomyosin represented as mechanical springs (blue and red represents junctional and medio-apical actomyosin, respectively). Under uniaxial tension, vinculin increases at the perpendicular junctions, indicating medio-apical springs generates the tension to recruit vinculin at those junctions.

Although vinculin recruitment is a well-established reporter of tension, previous studies have shown vinculin can also be recruited via force-independent mechanisms (Yonemura et al., 2010). To confirm that our results on medio-apical actomyosin represent transmitted forces upon stretching, we also performed laser ablations on horizontal junctions (hJC; parallel to the stretch axis), vertical junctions (vJC; perpendicular to the stretch axis), and medio-apical regions (MA) on relaxed and stretched tissues. We measured maximum ɛ_xx_ strain post-ablation to represent the regional tension (**Figure 5f**). In relaxed tissue, ablation-induced strain ɛ_xx_ was similar among hJC, vJC, and MA (**Figure 5f, and 5g**). However, under stretched conditions, ablation of the medio-apical region resulted in a significantly larger ɛ_xx_ strain than ablations at hJC and vJC (**Figure 5f, and 5g**). We find that the medio-apical domain of the cell experiences higher mechanical tension, consistent with vinculin localization; these findings indicate that medio-apical actomyosin serves as the primary force-transmitting structure during uniaxial stretch.

Together, these findings demonstrate that medio-apical actomyosin plays a dominant role in force transmission, whereas junctional actomyosin primarily facilitates force dissipation. Both vinculin recruitment and recoil after laser ablation during mechanical strain correlates with medio-apical tension rather than junctional strain (**Figure 5c-g, and 6b**), reinforcing the idea that epithelial force transmission occurs primarily through the medio-apical cytoskeletal network.

## 3. Conclusion and Discussion

In this study, we uncovered fundamental principles governing the heterogeneous mechanical behavior of epithelial tissues under uniaxial strain (**Figure 6a**). Individual cells within an epithelial tissue exhibit variable strain responses, with strain distribution correlating with cell surface area and cell intrinsic material properties. Previous studies have suggested heterogeneous mechanical behaviors in epithelial tissues; however, these insights were often derived from indirect measurements or computational models. For instance, computational modeling and force inference methods have been used to estimate intracellular tension and pressure differentials from tissue-level deformation in epithelial tissues (Sugimura and Ishihara, 2013; Etournay et al., 2015). While these approaches have provided valuable insights, they inferred mechanical properties rather than directly measured cellular strain under controlled strain conditions. With control over the mechanical environment, our study provides quantitative, single-cell level measurements of strain heterogeneity. This approach allows us to provide quantitative insights into the complex interplay between cellular architecture and force transmission in epithelial tissues, building upon and extending previous studies in the field of tissue mechanics (Gjorevski and Nelson, 2010; Thompson et al., 2019; Kim et al., 2020).

A key finding of our study, focusing on the Poisson effect, is the quantified contribution of intracellular microstructures. This effect is revealed in the strong correlation between anisotropy, defined by Poisson Index, and the ɛ_xx_ strain in individual cells. Cells with higher Poisson Index exhibited lower ɛ_xx_ strain. Furthermore, cells with larger compression along the apical-basal axis, represented by larger ɛ_zz_, exhibited higher ɛ_xx_ strain. This pattern suggests that epithelial cells behave as anisotropic materials, with mechanical properties that differ depending on the direction of applied force (**Figure 6a**). Such anisotropy likely arises from the complex internal architecture of epithelial cells, including the density and orientation of cytoskeletal elements and their localization to the apical cortex, and distribution of cellular organelles (Haase and Pelling, 2015; Bonakdar et al., 2016). Additionally, we observed weak, albeit statistically significant, correlation between cell surface area, cell orientation and strain. While the influences of cell size and cell orientation are less pronounced than that of the Poisson’s ratio, they still contribute to the overall mechanical behavior of the tissue and are also likely to reflect intracellular structures.

These geometric influences on cell responses to force add new dimensions to our understanding of epithelial mechanics, complementing previous work on the impact of cell shape on mechanosensing and mechanotransduction in tissues (McBeath et al., 2004; Lecuit and Lenne, 2007; Latorre et al., 2018). While we tested several geometric parameters, additional geometric features have been proposed to contribute to heterogeneous strain responses. For instance, force inference-based methods suggest that junctions with sharper angles experience higher stress (Brauns et al., 2024), raising the possibility that cells with acute-angled junctions might resist strain more efficiently. Future investigations are needed to test whether these geometric features contribute to strain heterogeneity in epithelial tissues.

Due to the prevailing thought that actomyosin levels and activity dictate mechanical properties of embryonic cells and tissues (for instance, see (Zhou et al., 2009; Chanet et al., 2017; Umetsu and Kuranaga, 2017; Miao and Blankenship, 2020; Kollimada et al., 2021)) we were surprised to find that initial pre-strain actomyosin levels did not contribute to the strain heterogeneity. Our results suggest that the dynamic regulation and spatial organization of actomyosin, rather than its absolute quantity, play a more critical role in determining a cell’s response to applied forces. While baseline actomyosin levels do not directly govern strain heterogeneity, actomyosin remains highly mechanoresponsive, as demonstrated in both our study and previous findings showing that actomyosin meshworks can sense and respond to mechanical cues by changing their conformation and orientation (Xie and Martin, 2015; Hara et al., 2016; Chanet et al., 2017; Sun et al., 2017).

Our investigation into actomyosin turn-over and stability uncovered distinct roles for junctional and medio-apical actomyosin pools in force transmission and dissipation. Notably, we find that medio-apical actomyosin serves as a primary player in force transmission, while junctional actomyosin appears more involved in force dissipation (**Figure 4, and 6b**). This finding contrasts with many traditional epithelial morphogenetic models that emphasize junctional forces (Roper, 2015; Staddon et al., 2019; Mira-Osuna and Borgne, 2024). Our findings align with and extend recent studies highlighting the crucial role of medio-apical actomyosin in epithelial mechanics. For instance, medio-apical contractile pulses have been shown to coordinate epithelial remodeling and drive apical constriction, reinforcing the idea that medio-apical actomyosin plays an essential role in tissue morphogenesis (Arnold et al., 2019; Rosa-Birriel et al., 2024). By emphasizing the importance of medio-apical actomyosin in force transmission, our study challenges the traditional focus on junctional forces and highlights the complex interplay between different actomyosin pools during epithelial morphogenesis (Jackson et al., 2017; Baldwin et al., 2023).

While our study provides important insights into the roles of junctional and medio-apical actomyosin in epithelial mechanics, several limitations should be considered. First, our stretcher design prevents us from investigating basal-lateral actomyosin contributions to force transmission and dissipation. Recent studies have shown that basal actomyosin networks play a crucial role in epithelial expansion and remodeling (Li et al., 2024). Additionally, our study primarily focused on responses immediately after application of strain; other relatively long-term dynamic mechanisms, such as cell division and radial intercalation, were not assessed. Externally applied tension has been shown to induce and re-orient cell division (Nestor-Bergmann et al., 2019). Other studies have shown that radial cell intercalation can significantly impact force distribution and tissue fluidity (Sedzinski et al., 2016) potentially altering the mechanical properties of the epithelium in ways not captured by our experimental design. Future studies are required to explore whether these dynamic processes contribute to force adaptation in epithelial tissues. Furthermore, our experimental setup did not account for the potential influence of other cytoskeleton components (e.g., intermediate filaments) and the extracellular matrix, which have both been shown to play a role in mechanotransduction (Sanghvi-Shah and Weber, 2017; Kozyrina et al., 2020; Duque et al., 2024).

The heterogeneous strain distribution observed in our study has important implications for tissue-level behaviors. Localized regions of high strain may serve an instructive cues gene induction and secondary patterning (Farge, 2003; Caldarelli et al., 2024). Sites of high strain may regulate morphogenesis through feedback (Sampathkumar et al., 2014; Gustafson et al., 2022), or trigger morphogenetic events such as wound healing (Berry et al., 2023), tissue folding, invagination (Lecuit and Lenne, 2007; Suzuki et al., 2024), regeneration (Vining and Mooney, 2017), or metastases (Northcott et al., 2018). Additionally, the ability of tissues to distribute strain non-uniformly may enhance their overall mechanical resilience, allowing them to withstand and adapt to a wider range of mechanical challenges. This principle of mechanical heterogeneity as a source of robustness may be a fundamental feature of epithelial tissue design, contributing to their remarkable ability to maintain integrity under diverse mechanical conditions (Hannezo and Heisenberg, 2019).

Our findings highlight key areas for further investigation, particularly in understanding how cells detect and respond to local strain variations within a tissue. Another critical aspect is uncovering the signaling pathways that regulate the differential activity of junctional and medio-apical actomyosin pools. Addressing these challenges will require a combination of advanced imaging techniques, computational models that capture the multi-scale complexity of tissue mechanics, and tools that can effectively and precisely manipulate tissue mechanics.

## 4. Materials and Methods

### Preparation of the TissueTractor system

Fabrication of the microscope stage insert and cassettes were previously described (Yang et al., 2025). In brief, all components of the microscope stage insert, except the picomotor piezo linear actuators (8301NF; Newport) and the 45 x 50 mm glass coverslip, were printed with Polylactic Acid (PLA; Prusa Research) filaments by a Fused Deposition Modeling 3D-printer (Prusa i3 MK3S; Prusa Research) (**Figure 1a**). The cassette was assembled using Polyester (PES) sheets (Red, Precision Brand), a Polydimethylsiloxane (PDMS) sheet (0.005’’, 40D, Gloss finish, Specialty Manufacturing, inc.), and two 3D-printed stretcher blocks (Clear, Form2, Formlabs) (**Figure 1a’**) (Yang et al., 2025).

### *Xenopus laevis* embryo culture, microinjection, and organotypic explant isolation and mounting

All *Xenopus laevis* work was approved by the University of Pittsburgh Division of Laboratory Animal Resources. *Xenopus laevis* embryos were obtained by the standard procedure and cultured in 1/3x modified Barth Solutions (MBS) (Sive et al., 2000) to desired stage (Nieuwkoop and Faber, 1967). Embryos were injected at either the 2-cell or the 4-cell stage. 200 pg of mem-mNeonGreen, mem-miRFP670, mem-BFP, or SF9-mNeonGreen was injected per embryo. 100 pg of LifeAct-mScarlet or LifeAct-eGFP was injected per embryo. 70 pg of Vinculin-mNeonGreen was injected per embryo.

Organotypic explants isolation and mounting on the cassette were previously described (Chen et al., 2024). In brief, cassettes were treated in an oxygen plasma cleaner (Harrick Plasma) for 2 minutes to activate the surface of the PDMS substrates and coated with 0.025 μg/μl of fibronectin (Chem Cruz) at room temperature for 1 hour or at 4°C overnight. Embryos at early gastrula stage (Stage 9-10) (Nieuwkoop and Faber, 1967) were transferred into Danilchik’s For Amy (DFA) (Sive et al., 2000) medium with antibiotic and antimycotic (Sigma). An organotypic animal cap explant was microsurgically removed (Joshi and Davidson, 2010) and positioned at the center of the PDMS substrate, immobilized by a glass coverslip bridge. The cassettes with the explants were incubated at 14°C to the desired stage (Yang et al., 2025).

### Stretching experiment setup and imaging

The cassette with the explant was transferred and positioned in the sample chamber of the stage insert with 4 mL DFA medium (Yang et al., 2025). Images were acquired at relaxed state (S_0_) and incrementally stretched, with images captured at each step (S_1_, S_2_, …, S_n_) while tracking the same group of cells throughout (**Figure 1b**) using an inverted compound microscope (leica) with a 25x/0.95NA water immersion objective lens, equipped with a spinning disk scanhead (Yokogawa) and a CMOS camera (Hamamatsu). Sequential images were acquired using a microscope automation software (μManager 2.0) (Edelstein et al., 2014). Images with fluorescent recovery after photobleaching were acquired on a confocal laser scanning microscope (SP5; Leica Microsystems) with a 25x water immersion objective lens with 2.5x zoom.

### Fluorescence recovery after photobleaching

Photobleaching was performed on selected regions of cell junctions or medio-apical surfaces with the 488 nm argon laser operating at 100% laser power for 3.6 seconds (1.2 second per frame for 3 frames). 10 Pre-bleaching images and 50 post-bleaching images were collected with 20% laser power. Image gain was turned up until F-actin in medio-apical region was visible to perform FRAP on the medio-apical surfaces of the cells. F-actin intensity was scaled using min-max normalization, with pre-bleach intensity as 1 and bleaching intensity as 0, and fitted using one phase decay in GraphPad Prism version 10.4.1 (GraphPad Software).

### Laser ablation

Laser ablation was performed with Micropoint system (Andor) using a nitrogen dye laser (435 nm) equipped in the spinning disk confocal microscope previously described. Micropoint was calibrated on a mirror slide for accuracy. The power of the laser is calibrated and tuned on a controlled explant before every experiment. A point ROI was selected for ablation either on cell junctions or on medio-apical surfaces of the cell. An image was taken before ablation and single z-slice images were taken every 200ms for 48 seconds in total.

### Image segmentation and analysis

Individual cells were segmented using an established segmentation tool SeedWater Segmenter (Mashburn et al., 2012), and a custom ImageJ macro (Table S1) was used to acquire cell ROIs and shape information from the segmented cells. A custom MATLAB m-code (Table S1) calculated individual cell engineering strains based on cell shape changes (Vijayraghavan, 2018; Yang et al., 2025). Cells undergo cell division were excluded from the analysis.

F-actin, myosin II intensities, and junctional vinculin intensities were normalized to membrane intensities of the same cell. Maximum ɛ_xx_ strain after laser ablation was calculated using a custom FIJI macro StrainMapper (Table S1) previously described (Stepien et al., 2019; Yang et al., 2025).

### Statistical Analysis

All statistical analysis was conducted in GraphPad Prism version 10.4.1 (GraphPad Software). Correlation and Pearson correlation coefficient were calculated by simple linear regression. Comparison for coordination numbers and vinculin intensity of different junctions were calculated using One Way ANOVA. FRAP recovery fraction was fitted using one phase decay. Half-life and immobile fraction, ɛ_xx_ strain after laser ablation were compared using Kruskal-Wallis test.

## Supplementary Materials

Supplementary materials are provided by the authors.

## Ethical Statement

All *Xenopus laevis* work was carried out according to Public Health Service (PHS) and United States Department of Agriculture (USDA) guidelines and approved by the Institutional Animal Care and Use Committee (IACUC) and the University of Pittsburgh Division of Laboratory Animal Resources (Protocol #24014521)

## Acknowledgement

We would like to thank Geneva Masak, Sommer Anjum, and other members of the Davidson group for their comments and discussions. This work was supported by the National Institutes of Health (R01 HD044750 and R37 HD044750). J.Y. was additionally supported by a Provost Dissertation Fellowship from the University of Pittsburgh. Y.W. and C.B.J. were additionally supported by Whiteford Faculty Funds and a Summer Undergraduate Research Internship (SURI) from the University of Pittsburgh Swanson School of Engineering. C.V.M. was additionally supported by a Brackenridge Fellowship from the University of Pittsburgh Frederick Honors College. We thank Dr. Ann Miller for sharing the *Xenopus laevis* Vinculin-eGFP and mem-mTagBFP2 plasmids and Drs. Ann Miller and Dr. Edwin Munro for sharing the SF9 myosin II activity reporter.

## References

Anjum, S., Turner, L., Atieh, Y., Eisenhoffer, G. T. and Davidson, L. A. (2024) ‘Assessing mechanical agency during apical apoptotic cell extrusion’, iScience 27(11): 111017.

Arnold, T. R., Shawky, J. H., Stephenson, R. E., Dinshaw, K. M., Higashi, T., Huq, F., Davidson, L. A. and Miller, A. L. (2019) ‘Anillin regulates epithelial cell mechanics by structuring the medial-apical actomyosin network’, Elife 8.

Atia, L., Fredberg, J. J., Gov, N. S. and Pegoraro, A. F. (2021) ‘Are cell jamming and unjamming essential in tissue development?’, Cells Dev 168: 203727.

Bachir, A. I., Horwitz, A. R., Nelson, W. J. and Bianchini, J. M. (2017) ‘Actin-Based Adhesion Modules Mediate Cell Interactions with the Extracellular Matrix and Neighboring Cells’, Cold Spring Harb Perspect Biol 9(7).

Baldwin, A., Popov, I. K., Keller, R., Wallingford, J. and Chang, C. (2023) ‘The RhoGEF protein Plekhg5 regulates medioapical and junctional actomyosin dynamics of apical constriction during Xenopus gastrulation’, Mol Biol Cell 34(7): ar64.

Banavar, S. P., Carn, E. K., Rowghanian, P., Stooke-Vaughan, G., Kim, S. and Campas, O. (2021) ‘Mechanical control of tissue shape and morphogenetic flows during vertebrate body axis elongation’, Sci Rep 11(1): 8591.

Barriga, E. H., Franze, K., Charras, G. and Mayor, R. (2018) ‘Tissue stiffening coordinates morphogenesis by triggering collective cell migration in vivo’, Nature 554(7693): 523–527.

Benink, H. A. and Bement, W. M. (2005) ‘Concentric zones of active RhoA and Cdc42 around single cell wounds’, J Cell Biol 168(3): 429–39.

Berry, C. E., Downer, M., Jr., Morgan, A. G., Griffin, M., Liang, N. E., Kameni, L., Laufey Parker, J. B., Guo, J., Longaker, M. T. and Wan, D. C. (2023) ‘The effects of mechanical force on fibroblast behavior in cutaneous injury’, Front Surg 10: 1167067.

Bi, D., Yang, X., Marchetti, M. C. and Manning, M. L. (2016) ‘Motility-driven glass and jamming transitions in biological tissues’, Phys Rev X 6(2).

Blauth, E., Kubitschke, H., Gottheil, P., Grosser, S. and Käs, J. A. A. (2021) ‘Jamming in Embryogenesis and Cancer Progression’, Frontiers in Physics 9.

Bodenschatz, J. F. E., Ajmail, K., Skamrahl, M., Vache, M., Gottwald, J., Nehls, S. and Janshoff, A. (2022) ‘Epithelial cells sacrifice excess area to preserve fluidity in response to external mechanical stress’, Commun Biol 5(1): 855.

Bonakdar, N., Gerum, R., Kuhn, M., Sporrer, M., Lippert, A., Schneider, W., Aifantis, K. E. and Fabry, B. (2016) ‘Mechanical plasticity of cells’, Nat Mater 15(10): 1090–4.

Brauns, F., Claussen, N. H., Lefebvre, M. F., Wieschaus, E. F. and Shraiman, B. I. (2024) ‘The geometric basis of epithelial convergent extension’, Elife 13.

Caldarelli, P., Chamolly, A., Villedieu, A., Alegria-Prevot, O., Phan, C., Gros, J. and Corson, F. (2024) ‘Self-organized tissue mechanics underlie embryonic regulation’, Nature 633(8031): 887–894.

Chanet, S., Miller, C. J., Vaishnav, E. D., Ermentrout, B., Davidson, L. A. and Martin, A. C. (2017) ‘Actomyosin meshwork mechanosensing enables tissue shape to orient cell force’, Nat Commun 8: 15014.

Chang, A. Y. and Marshall, W. F. (2017) ‘Organelles - understanding noise and heterogeneity in cell biology at an intermediate scale’, J Cell Sci 130(5): 819–826.

Chen, Y., Li, Z., Kong, F., Ju, L. A. and Zhu, C. (2024) ‘Force-Regulated Spontaneous Conformational Changes of Integrins alpha(5)beta(1) and alpha(V)beta(3)’, ACS Nano 18(1): 299–313.

Chien, Y. H., Keller, R., Kintner, C. and Shook, D. R. (2015) ‘Mechanical strain determines the axis of planar polarity in ciliated epithelia’, Curr Biol 25(21): 2774–2784.

Chirasani, V. R., Khan, M. A. I., Malavade, J. N., Dokholyan, N. V., Hoffman, B. D. and Campbell, S. L. (2023) ‘Molecular basis and cellular functions of vinculin-actin directional catch bonding’, Nat Commun 14(1): 8300.

Chu, C., Masak, G., Yang, J. and Davidson, L. (2020) ‘From biomechanics to mechanobiology: Xenopus provides direct access to the physical principles that shape the embryo’, Current Opinion in Genetics & Development 63: 71–77.

Clarke, D. N. and Martin, A. C. (2021) ‘Actin-based force generation and cell adhesion in tissue morphogenesis’, Curr Biol 31(10): R667–R680.

Collinet, C. and Lecuit, T. (2013) ‘Stability and dynamics of cell-cell junctions’, Prog Mol Biol Transl Sci 116: 25–47.

Conway, J. R. W., Isomursu, A., Follain, G., Harma, V., Jou-Olle, E., Pasquier, N., Valimaki, E. P. O., Rantala, J. K. and Ivaska, J. (2023) ‘Defined extracellular matrix compositions support stiffness-insensitive cell spreading and adhesion signaling’, Proc Natl Acad Sci U S A 120(43): e2304288120.

Crossley, R. M., Johnson, S., Tsingos, E., Bell, Z., Berardi, M., Botticelli, M., Braat, Q. J. S., Metzcar, J., Ruscone, M., Yin, Y. et al. (2024) ‘Modeling the extracellular matrix in cell migration and morphogenesis: a guide for the curious biologist’, Front Cell Dev Biol 12: 1354132.

Curran, S., Strandkvist, C., Bathmann, J., de Gennes, M., Kabla, A., Salbreux, G. and Baum, B. (2017) ‘Myosin II Controls Junction Fluctuations to Guide Epithelial Tissue Ordering’, Dev Cell 43(4): 480–492 e6.

Davidson, L., von Dassow, M. and Zhou, J. (2009) ‘Multi-scale mechanics from molecules to morphogenesis’, Int J Biochem Cell Biol 41(11): 2147–62.

Duque, J., Bonfanti, A., Fouchard, J., Baldauf, L., Azenha, S. R., Ferber, E., Harris, A., Barriga, E. H., Kabla, A. J. and Charras, G. (2024) ‘Rupture strength of living cell monolayers’, Nat Mater 23(11): 1563–1574.

Eckert, J., Ladoux, B., Mege, R. M., Giomi, L. and Schmidt, T. (2023) ‘Hexanematic crossover in epithelial monolayers depends on cell adhesion and cell density’, Nat Commun 14(1): 5762.

Edelstein, A. D., Tsuchida, M. A., Amodaj, N., Pinkard, H., Vale, R. D. and Stuurman, N. (2014) ‘Advanced methods of microscope control using muManager software’, J Biol Methods 1(2).

Engler, A. J., Sen, S., Sweeney, H. L. and Discher, D. E. (2006) ‘Matrix elasticity directs stem cell lineage specification’, Cell 126(4): 677–89.

Erdemci-Tandogan, G., Clark, M. J., Amack, J. D. and Manning, M. L. (2018) ‘Tissue Flow Induces Cell Shape Changes During Organogenesis’, Biophys J 115(11): 2259–2270.

Etournay, R., Popovic, M., Merkel, M., Nandi, A., Blasse, C., Aigouy, B., Brandl, H., Myers, G., Salbreux, G., Julicher, F. et al. (2015) ‘Interplay of cell dynamics and epithelial tension during morphogenesis of the Drosophila pupal wing’, Elife 4: e07090.

Farge, E. (2003) ‘Mechanical induction of Twist in the Drosophila foregut/stomodeal primordium’, Curr Biol 13(16): 1365–77.

Feroze, R., Shawky, J. H., von Dassow, M. and Davidson, L. A. (2015) ‘Mechanics of blastopore closure during amphibian gastrulation’, Dev Biol 398(1): 57–67.

Founounou, N., Farhadifar, R., Collu, G. M., Weber, U., Shelley, M. J. and Mlodzik, M. (2021) ‘Tissue fluidity mediated by adherens junction dynamics promotes planar cell polarity-driven ommatidial rotation’, Nat Commun 12(1): 6974.

Gibson, M. C., Patel, A. B., Nagpal, R. and Perrimon, N. (2006) ‘The emergence of geometric order in proliferating metazoan epithelia’, Nature 442(7106): 1038–41.

Gjorevski, N. and Nelson, C. M. (2010) ‘The mechanics of development: Models and methods for tissue morphogenesis’, Birth Defects Res C Embryo Today 90(3): 193–202.

Gjorevski, N. and Nelson, C. M. (2012) ‘Mapping of mechanical strains and stresses around quiescent engineered three-dimensional epithelial tissues’, Biophys J 103(1): 152–62.

Gomez, G. A., McLachlan, R. W. and Yap, A. S. (2011) ‘Productive tension: force-sensing and homeostasis of cell-cell junctions’, Trends Cell Biol 21(9): 499–505.

Greaves, G. N., Greer, A. L., Lakes, R. S. and Rouxel, T. (2011) ’Poisson’s ratio and modern materials’, Nat Mater 10(11): 823–37.

Guillot, C. and Lecuit, T. (2013) ‘Mechanics of epithelial tissue homeostasis and morphogenesis’, Science 340(6137): 1185–9.

Guo, M., Ehrlicher, A. J., Jensen, M. H., Renz, M., Moore, J. R., Goldman, R. D., Lippincott-Schwartz, J., Mackintosh, F. C. and Weitz, D. A. (2014) ‘Probing the stochastic, motor-driven properties of the cytoplasm using force spectrum microscopy’, Cell 158(4): 822–832.

Gustafson, H. J., Claussen, N., De Renzis, S. and Streichan, S. J. (2022) ‘Patterned mechanical feedback establishes a global myosin gradient’, Nat Commun 13(1): 7050.

Haase, K. and Pelling, A. E. (2015) ‘Investigating cell mechanics with atomic force microscopy’, J R Soc Interface 12(104): 20140970.

Hannezo, E. and Heisenberg, C. P. (2019) ‘Mechanochemical Feedback Loops in Development and Disease’, Cell 178(1): 12–25.

Hara, Y., Shagirov, M. and Toyama, Y. (2016) ‘Cell Boundary Elongation by Non-autonomous Contractility in Cell Oscillation’, Curr Biol 26(17): 2388–96.

Heer, N. C. and Martin, A. C. (2017) ‘Tension, contraction and tissue morphogenesis’, Development 144(23): 4249–4260.

Hidalgo-Carcedo, C., Hooper, S., Chaudhry, S. I., Williamson, P., Harrington, K., Leitinger, B. and Sahai, E. (2011) ‘Collective cell migration requires suppression of actomyosin at cell-cell contacts mediated by DDR1 and the cell polarity regulators Par3 and Par6’, Nat Cell Biol 13(1): 49–58.

Holle, A. W., Young, J. L., Van Vliet, K. J., Kamm, R. D., Discher, D., Janmey, P., Spatz, J. P. and Saif, T. (2018) ‘Cell-Extracellular Matrix Mechanobiology: Forceful Tools and Emerging Needs for Basic and Translational Research’, Nano Lett 18(1): 1–8.

Huebner, R. J., Malmi-Kakkada, A. N., Sarikaya, S., Weng, S., Thirumalai, D. and Wallingford, J. B. (2021) ‘Mechanical heterogeneity along single cell-cell junctions is driven by lateral clustering of cadherins during vertebrate axis elongation’, Elife 10.

Ingber, D. E., Prusty, D., Sun, Z., Betensky, H. and Wang, N. (1995) ‘Cell shape, cytoskeletal mechanics, and cell cycle control in angiogenesis’, J Biomech 28(12): 1471–84.

Jackson, T. R., Kim, H. Y., Balakrishnan, U. L., Stuckenholz, C. and Davidson, L. A. (2017) ‘Spatiotemporally Controlled Mechanical Cues Drive Progenitor Mesenchymal-to-Epithelial Transition Enabling Proper Heart Formation and Function’, Curr Biol 27(9): 1326–1335.

Jain, A., Ulman, V., Mukherjee, A., Prakash, M., Cuenca, M. B., Pimpale, L. G., Munster, S., Haase, R., Panfilio, K. A., Jug, F. et al. (2020) ‘Regionalized tissue fluidization is required for epithelial gap closure during insect gastrulation’, Nat Commun 11(1): 5604.

Joshi, S. D. and Davidson, L. A. (2010) ‘Live-cell imaging and quantitative analysis of embryonic epithelial cells in Xenopus laevis’, J Vis Exp(39).

Jung, A. C., Ribeiro, C., Michaut, L., Certa, U. and Affolter, M. (2006) ‘Polychaetoid/ZO-1 is required for cell specification and rearrangement during Drosophila tracheal morphogenesis’, Curr Biol 16(12): 1224–31.

Kale, G. R., Yang, X., Philippe, J. M., Mani, M., Lenne, P. F. and Lecuit, T. (2018) ‘Distinct contributions of tensile and shear stress on E-cadherin levels during morphogenesis’, Nat Commun 9(1): 5021.

Khalilgharibi, N., Fouchard, J., Asadipour, N., Barrientos, R., Duda, M., Bonfanti, A., Yonis, A., Harris, A., Mosaffa, P., Fujita, Y. et al. (2019) ‘Stress relaxation in epithelial monolayers is controlled by the actomyosin cortex’, Nature Physics 15(8): 839–847.

Kim, H. Y., Jackson, T. R., Stuckenholz, C. and Davidson, L. A. (2020) ‘Tissue mechanics drives regeneration of a mucociliated epidermis on the surface of Xenopus embryonic aggregates’, Nat Commun 11(1): 665.

Kollimada, S., Senger, F., Vignaud, T., Thery, M., Blanchoin, L. and Kurzawa, L. (2021) ‘The biochemical composition of the actomyosin network sets the magnitude of cellular traction forces’, Mol Biol Cell 32(18): 1737–1748.

Kowalczyk, A. P. and Green, K. J. (2013) ‘Structure, function, and regulation of desmosomes’, Prog Mol Biol Transl Sci 116: 95–118.

Kozyrina, A. N., Piskova, T. and Di Russo, J. (2020) ‘Mechanobiology of Epithelia From the Perspective of Extracellular Matrix Heterogeneity’, Front Bioeng Biotechnol 8: 596599.

Landino, J., Misterovich, E., van den Goor, L., Adhikary, B., Chumki, S., Davidson, L. A. and Miller, A. L. (2025) ‘Neighbor cells restrain furrowing during Xenopus epithelial cytokinesis’, Dev Cell.

Latorre, E., Kale, S., Casares, L., Gomez-Gonzalez, M., Uroz, M., Valon, L., Nair, R. V., Garreta, E., Montserrat, N., Del Campo, A. et al. (2018) ‘Active superelasticity in three-dimensional epithelia of controlled shape’, Nature 563(7730): 203–208.

Lecuit, T. and Lenne, P. F. (2007) ‘Cell surface mechanics and the control of cell shape, tissue patterns and morphogenesis’, Nat Rev Mol Cell Biol 8(8): 633–44.

Li, S., Liu, Z. Y., Li, H., Zhou, S., Liu, J., Sun, N., Yang, K. F., Dougados, V., Mangeat, T., Belguise, K. et al. (2024) ‘Basal actomyosin pulses expand epithelium coordinating cell flattening and tissue elongation’, Nat Commun 15(1): 3000.

Li, X., Das, A. and Bi, D. (2019) ‘Mechanical Heterogeneity in Tissues Promotes Rigidity and Controls Cellular Invasion’, Phys Rev Lett 123(5): 058101.

Li, Y., Huang, G., Li, M., Wang, L., Elson, E. L., Lu, T. J., Genin, G. M. and Xu, F. (2016) ‘An approach to quantifying 3D responses of cells to extreme strain’, Sci Rep 6: 19550.

Luby-Phelps, K. (2000) ‘Cytoarchitecture and physical properties of cytoplasm: volume, viscosity, diffusion, intracellular surface area’, Int Rev Cytol 192: 189–221.

Maller, O., Martinson, H. and Schedin, P. (2010) ‘Extracellular matrix composition reveals complex and dynamic stromal-epithelial interactions in the mammary gland’, J Mammary Gland Biol Neoplasia 15(3): 301–18.

Manning, L. A., Perez-Vale, K. Z., Schaefer, K. N., Sewell, M. T. and Peifer, M. (2019) ‘The Drosophila Afadin and ZO-1 homologues Canoe and Polychaetoid act in parallel to maintain epithelial integrity when challenged by adherens junction remodeling’, Mol Biol Cell 30(16): 1938–1960.

Mashburn, D. N., Lynch, H. E., Ma, X. and Hutson, M. S. (2012) ‘Enabling user-guided segmentation and tracking of surface-labeled cells in time-lapse image sets of living tissues’, Cytometry A 81(5): 409–18.

McBeath, R., Pirone, D. M., Nelson, C. M., Bhadriraju, K. and Chen, C. S. (2004) ‘Cell shape, cytoskeletal tension, and RhoA regulate stem cell lineage commitment’, Dev Cell 6(4): 483–95.

Miao, H. and Blankenship, J. T. (2020) ‘The pulse of morphogenesis: actomyosin dynamics and regulation in epithelia’, Development 147(17).

Mira-Osuna, M. and Borgne, R. L. (2024) ‘Assembly, dynamics and remodeling of epithelial cell junctions throughout development’, Development 151(1).

Munjal, A. and Lecuit, T. (2014) ‘Actomyosin networks and tissue morphogenesis’, Development 141(9): 1789–93.

Ndiaye, A. B., Koenderink, G. H. and Shemesh, M. (2022) ‘Intermediate Filaments in Cellular Mechanoresponsiveness: Mediating Cytoskeletal Crosstalk From Membrane to Nucleus and Back’, Front Cell Dev Biol 10: 882037.

Nestor-Bergmann, A., Blanchard, G. B., Hervieux, N., Fletcher, A. G., Etienne, J. and Sanson, B. (2022) ‘Adhesion-regulated junction slippage controls cell intercalation dynamics in an Apposed-Cortex Adhesion Model’, PLoS Comput Biol 18(1): e1009812.

Nestor-Bergmann, A., Stooke-Vaughan, G. A., Goddard, G. K., Starborg, T., Jensen, O. E. and Woolner, S. (2019) ‘Decoupling the Roles of Cell Shape and Mechanical Stress in Orienting and Cueing Epithelial Mitosis’, Cell Rep 26(8): 2088–2100 e4.

Nieuwkoop, P. D. and Faber, J. (1967) Normal tables of Xenopus laevis (Daudin): Amsterdam: Elsevier North-Holland Biomedical Press.

Nizak, C., Martin-Lluesma, S., Moutel, S., Roux, A., Kreis, T. E., Goud, B. and Perez, F. (2003) ‘Recombinant antibodies against subcellular fractions used to track endogenous Golgi protein dynamics in vivo’, Traffic 4(11): 739–53.

Northcott, J. M., Dean, I. S., Mouw, J. K. and Weaver, V. M. (2018) ‘Feeling Stress: The Mechanics of Cancer Progression and Aggression’, Front Cell Dev Biol 6: 17.

O’Keeffe, M (1979) ‘A proposed rigorous definition of coordination number’, Foundations of Crystallography 35(5): 772–775.

Oakes, P. W., Banerjee, S., Marchetti, M. C. and Gardel, M. L. (2014) ‘Geometry regulates traction stresses in adherent cells’, Biophys J 107(4): 825–33.

Palmquist, Karl H, Tiemann, Sydney F, Ezzeddine, Farrah L, Yang, Sichen, Pfeifer, Charlotte R, Erzberger, Anna, Rodrigues, Alan R and Shyer, Amy E (2022) ‘Reciprocal cell-ECM dynamics generate supracellular fluidity underlying spontaneous follicle patterning’, Cell 185(11): 1960–1973. e11.

Quiros, M. and Nusrat, A. (2014) ‘RhoGTPases, actomyosin signaling and regulation of the epithelial Apical Junctional Complex’, Semin Cell Dev Biol 36: 194–203.

Roper, K. (2015) ‘Integration of cell-cell adhesion and contractile actomyosin activity during morphogenesis’, Curr Top Dev Biol 112: 103–27.

Rosa-Birriel, C., Malin, J. and Hatini, V. (2024) ‘Medioapical contractile pulses coordinated between cells regulate Drosophila eye morphogenesis’, J Cell Biol 223(2).

Ruppel, A., Worthmuller, D., Misiak, V., Kelkar, M., Wang, I., Moreau, P., Mery, A., Revilloud, J., Charras, G., Cappello, G. et al. (2023) ‘Force propagation between epithelial cells depends on active coupling and mechano-structural polarization’, Elife 12.

Sala, Stefano, Caillier, Alexia and Oakes, Patrick W (2024) ‘Principles and regulation of mechanosensing’, Journal of cell science 137(18).

Sampathkumar, A., Yan, A., Krupinski, P. and Meyerowitz, E. M. (2014) ‘Physical forces regulate plant development and morphogenesis’, Curr Biol 24(10): R475–83.

Sanghvi-Shah, R. and Weber, G. F. (2017) ‘Intermediate Filaments at the Junction of Mechanotransduction, Migration, and Development’, Front Cell Dev Biol 5: 81.

Saraswathibhatla, A., Indana, D. and Chaudhuri, O. (2023) ‘Cell-extracellular matrix mechanotransduction in 3D’, Nat Rev Mol Cell Biol 24(7): 495–516.

Sedzinski, J., Hannezo, E., Tu, F., Biro, M. and Wallingford, J. B. (2016) ‘Emergence of an Apical Epithelial Cell Surface In Vivo’, Dev Cell 36(1): 24–35.

Shawky, J. H., Balakrishnan, U. L., Stuckenholz, C. and Davidson, L. A. (2018) ‘Multiscale analysis of architecture, cell size and the cell cortex reveals cortical F-actin density and composition are major contributors to mechanical properties during convergent extension’, Development 145(19).

Shawky, J. H. and Davidson, L. A. (2015) ‘Tissue mechanics and adhesion during embryo development’, Dev Biol 401(1): 152–64.

Shellard, A. and Mayor, R. (2021) ‘Collective durotaxis along a self-generated stiffness gradient in vivo’, Nature 600(7890): 690–694.

Shyer, A. E., Rodrigues, A. R., Schroeder, G. G., Kassianidou, E., Kumar, S. and Harland, R. M. (2017) ‘Emergent cellular self-organization and mechanosensation initiate follicle pattern in the avian skin’, Science 357(6353): 811–815.

Sive, Hazel L., Grainger, Robert M. and Harland, Richard M. (2000) Early development of Xenopus laevis : a laboratory manual, Cold Spring Harbor, N.Y.: Cold Spring Harbor Laboratory Press.

Sprague, B. L. and McNally, J. G. (2005) ‘FRAP analysis of binding: proper and fitting’, Trends Cell Biol 15(2): 84–91.

Staddon, M. F., Cavanaugh, K. E., Munro, E. M., Gardel, M. L. and Banerjee, S. (2019) ‘Mechanosensitive Junction Remodeling Promotes Robust Epithelial Morphogenesis’, Biophys J 117(9): 1739–1750.

Stepien, T. L., Lynch, H. E., Yancey, S. X., Dempsey, L. and Davidson, L. A. (2019) ‘Using a continuum model to decipher the mechanics of embryonic tissue spreading from time-lapse image sequences: An approximate Bayesian computation approach’, PLoS One 14(6): 460774.

Sugimura, K. and Ishihara, S. (2013) ‘The mechanical anisotropy in a tissue promotes ordering in hexagonal cell packing’, Development 140(19): 4091–101.

Sumi, A., Hayes, P., D’Angelo, A., Colombelli, J., Salbreux, G., Dierkes, K. and Solon, J. (2018) ’Adherens Junction Length during Tissue Contraction Is Controlled by the Mechanosensitive Activity of Actomyosin and Junctional Recycling’, Dev Cell 47(4): 453–463 e3.

Sun, Z., Amourda, C., Shagirov, M., Hara, Y., Saunders, T. E. and Toyama, Y. (2017) ‘Basolateral protrusion and apical contraction cooperatively drive Drosophila germ-band extension’, Nat Cell Biol 19(4): 375–383.

Suzuki, M., Yasue, N. and Ueno, N. (2024) ‘Differential cellular stiffness across tissues that contribute to Xenopus neural tube closure’, Dev Growth Differ 66(5): 320–328.

Tarannum, Nawseen, Hargreaves, Dionn, Ilieva, Dessislava, Goddard, Georgina K., Jensen, Oliver E. and Woolner, Sarah (2024) ‘Mitotic spindle orientation and dynamics are fine-tuned by anisotropic tissue stretch via NuMA localisation’, bioRxiv.

Thompson, A. J., Pillai, E. K., Dimov, I. B., Foster, S. K., Holt, C. E. and Franze, K. (2019) ‘Rapid changes in tissue mechanics regulate cell behaviour in the developing embryonic brain’, Elife 8.

Tsuboi, A., Umetsu, D., Kuranaga, E. and Fujimoto, K. (2017) ‘Inference of Cell Mechanics in Heterogeneous Epithelial Tissue Based on Multivariate Clone Shape Quantification’, Front Cell Dev Biol 5: 68.

Umetsu, D. and Kuranaga, E. (2017) ‘Planar polarized contractile actomyosin networks in dynamic tissue morphogenesis’, Curr Opin Genet Dev 45: 90–96.

van Bodegraven, E. J. and Etienne-Manneville, S. (2021) ‘Intermediate Filaments from Tissue Integrity to Single Molecule Mechanics’, Cells 10(8).

Vasquez, C. G. and Martin, A. C. (2016) ‘Force transmission in epithelial tissues’, Dev Dyn 245(3): 361–71.

Vijayraghavan, D. S. and Davidson, L. A. (2017) ‘Mechanics of neurulation: From classical to current perspectives on the physical mechanics that shape, fold, and form the neural tube’, Birth Defects Res 109(2): 153–168.

Vijayraghavan, Deepthi (2018) Elucidating the Role of Mechanics in Neural Plate Convergent Extension Bioengineering., vol. PhD. Pittsburgh, PA: University of Pittsburgh.

Vining, K. H. and Mooney, D. J. (2017) ‘Mechanical forces direct stem cell behaviour in development and regeneration’, Nat Rev Mol Cell Biol 18(12): 728–742.

Xie, S. and Martin, A. C. (2015) ‘Intracellular signalling and intercellular coupling coordinate heterogeneous contractile events to facilitate tissue folding’, Nat Commun 6: 7161.

Yang, J., Hearty, E., Wang, Y., Vijayraghavan, D. S., Walter, T., Anjum, S., Stuckenholz, C., Cheng, Y. W., Balasubramanian, S., Dong, Y. et al. (2025) ‘The TissueTractor: A Device for Applying Large Strains to Tissues and Cells for Simultaneous High-Resolution Live Cell Microscopy’, Small Methods: e2500136.

Yonemura, S., Wada, Y., Watanabe, T., Nagafuchi, A. and Shibata, M. (2010) ‘alpha-Catenin as a tension transducer that induces adherens junction development’, Nat Cell Biol 12(6): 533–42.

Zhang, Y., He, Y., Bharadwaj, S., Hammam, N., Carnagey, K., Myers, R., Atala, A. and Van Dyke, M. (2009) ‘Tissue-specific extracellular matrix coatings for the promotion of cell proliferation and maintenance of cell phenotype’, Biomaterials 30(23-24): 4021–8.

Zhou, J., Kim, H. Y. and Davidson, L. A. (2009) ‘Actomyosin stiffens the vertebrate embryo during crucial stages of elongation and neural tube closure’, Development 136(4): 677–88.

Zhou, J., Kim, H. Y., Wang, J. H. and Davidson, L. A. (2010) ‘Macroscopic stiffening of embryonic tissues via microtubules, RhoGEF and the assembly of contractile bundles of actomyosin’, Development 137(16): 2785–94.

Zhou, J., Pal, S., Maiti, S. and Davidson, L. A. (2015) ‘Force production and mechanical accommodation during convergent extension’, Development 142(4): 692–701.

