## Supplementary Materials for "Shape, Strain, and Stability: Epithelia Under High Strain"

**Figure S1**

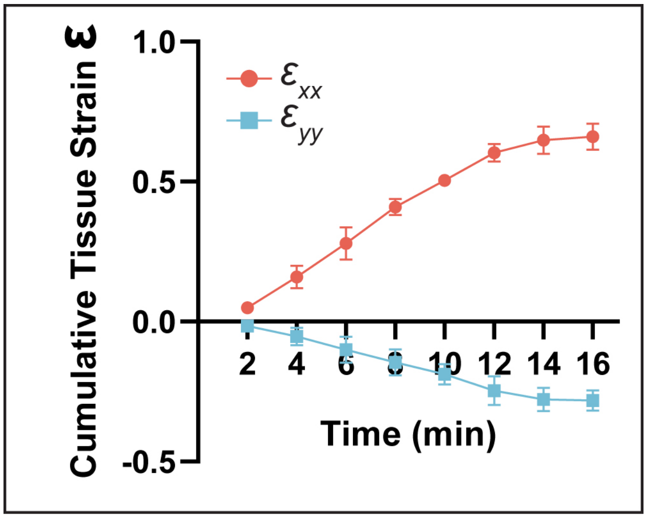


**Figure S1. Tissue Strain $\varepsilon_{xx}$ and $\varepsilon_{yy}$ at each stretch step (2 min, 4 min, …, 16 min).** Error bars, standard deviation. N = 3 tissues.

**Figure S2**

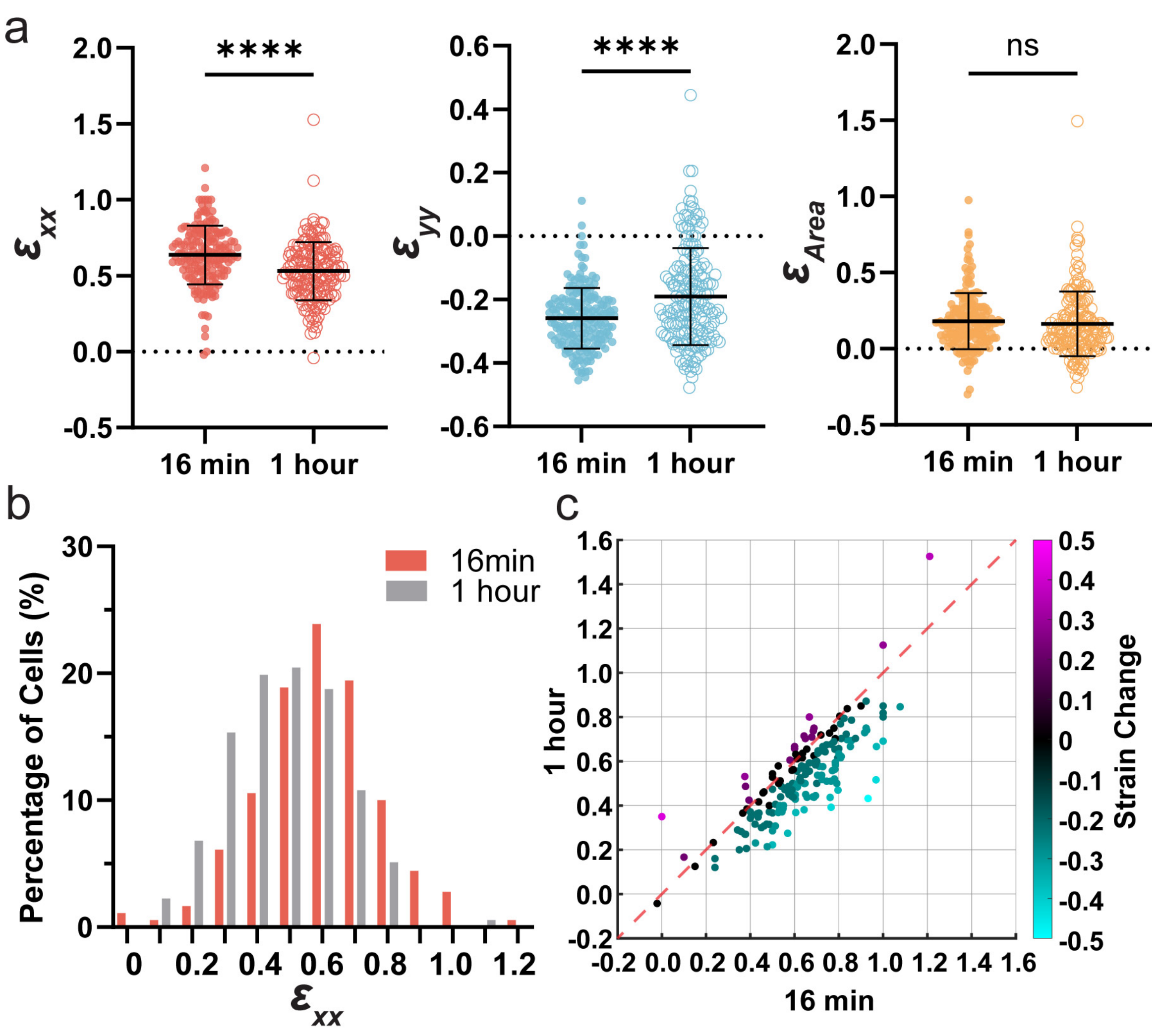


**Figure S2. Comparisons of cellular strains at maximum stretched position at 16 min and at 1 hour indicate relaxation of individual cells.** (**a**) Cellular strain $\varepsilon_{xx}$, $\varepsilon_{yy}$, and $\varepsilon_{Area}$ at 16 min and at 1 hour. Both $\varepsilon_{xx}$, $\varepsilon_{yy}$ show significant decrease, indicating relaxation of the cells, with $\varepsilon_{Area}$ remains the same. Error bars, standard deviation. (**b**) Frequency distribution of cells in binned $\varepsilon_{xx}$ strain at 16 min and at 1 hour. Individual bin size is 0.1. (c) Individual cellular strain $\varepsilon_{xx}$ plotted at 16 min against their corresponding values at 1 hour. The calibration bar shows strain change, with magenta indicating increase and cyan indicating decrease in strain $\varepsilon_{xx}$, and black indicating no change in strain between 16 min and 1 hour. The red dashed line represents the line of identity, where strain at 1 hour equals strain at 16 min. N = 181 cells.

**Figure S3**

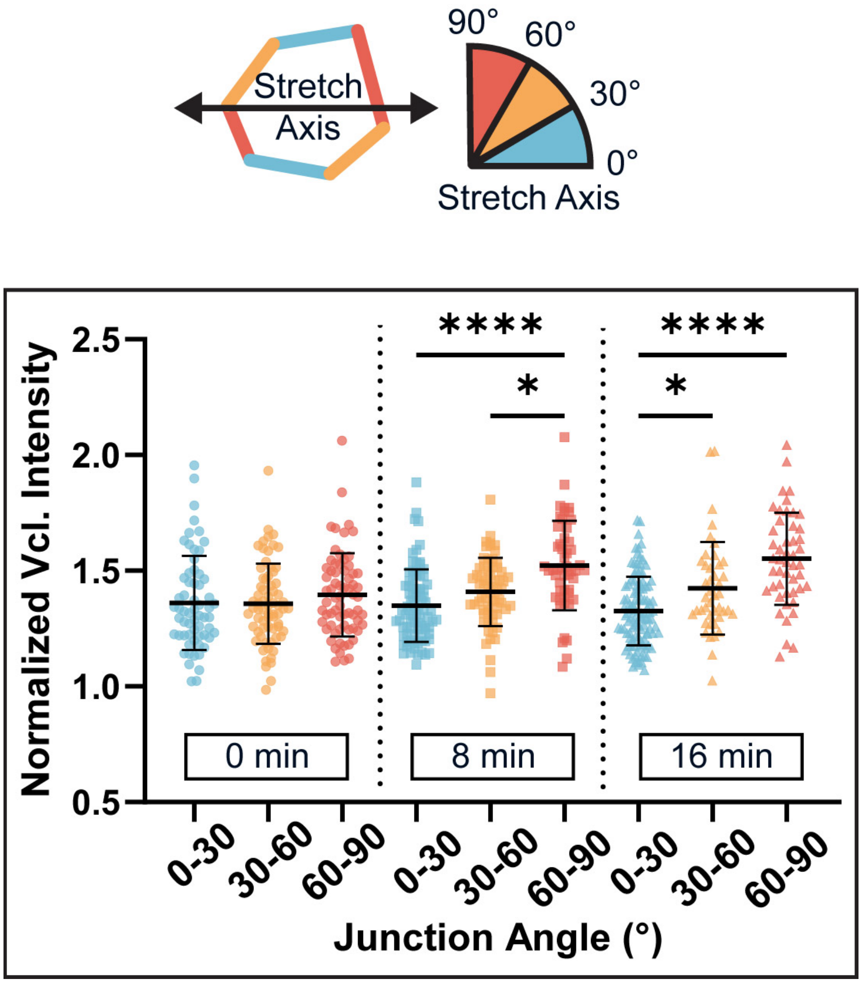


**Figure S3. Vinculin recruitment increases in junctions oriented 30° - 60° and 60° - 90° in stretched tissues.** A schematic illustrates cell junctions at three angle categories respective to the stretch axis: 0° - 30° (blue), 30° - 60° (yellow), and 60° - 90° (red). Quantifications of normalized vinculin intensity from three groups at 0 min (relaxed), 8 min, and 16 min). N = 199 junctions. Error bars, standard deviations. *Data shown are from the same dataset as in **Figure 5d**, reorganized to show clearer in-frame comparison across time points.

**Table S1. Image Analysis Macros**

| Macro Name | Author | Description | Input File Types | Input Parameters | Macro Category | Output Data | Output File Types | Software |
|---|---|---|---|---|---|---|---|---|
| Cell Descriptor | Deepthi Vijayraghavan | Generates FIJI ROI and assign cell IDs for each cell following SeedWater Segmenter. | Tiff | | Image Segmentation | Cell ROI | zip | FIJI |
| InfoForCellStrainBox_Perimeter | Jing Yang | Measures cell geometries including area, perimeter, orientation, intensity, width, height, slice number. | Tiff, zip | Cell ROI | Geometry Analysis | Cell area, perimeter, orientation, intensity, width, height, slice number | csv | FIJI |
| CellBoxStrain | Jing Yang, Deepthi Vijayraghavan | Calculates individual cellular strain rate and cumulative strain. | csv | Slice number | Strain Analysis | Cellular strain rate and cumulative strain | csv | MATLAB |
| ColorCodeMatlabStrain | Jing Yang | Color Code individual cell with strain calculated from the MATLAB analysis. | csv | Stack width, stack height, slice number | Image Generation | Color coded cells with respective cellular strain | Tiff | FIJI |
| AddROIsCreateTissue | Jing Yang | Add cell ROIs in the field to create tissue outline. | Tiff, zip | Cell ROI | Image Generation | Tissue ROI | zip | FIJI |
